# ComplexBrowser: a tool for identification and quantification of protein complexes in large scale proteomics datasets

**DOI:** 10.1101/573774

**Authors:** Wojciech Michalak, Vasileios Tsiamis, Veit Schwämmle, Adelina Rogowska-Wrzesińska

## Abstract

We have developed ComplexBrowser, an open source, online platform for supervised analysis of quantitative proteomics data that focuses on protein complexes. The software uses information from CORUM and Complex Portal databases to identify protein complex components. Based on the expression changes of individual complex subunits across the proteomics experiment it calculates Complex Fold Change (CFC) factor that characterises the overall protein complex expression trend and the level of subunit co-regulation. Thus up- and down-regulated complexes can be identified. It provides interactive visualisation of protein complexes composition and expression for exploratory analysis. It also incorporates a quality control step that includes normalisation and statistical analysis based on Limma test. ComplexBrowser performance was tested on two previously published proteomics studies identifying changes in protein expression in human adenocarcinoma tissue and during activation of mouse T-cells. The analysis revealed 1519 and 332 protein complexes, of which 233 and 41 were found co-ordinately regulated in the respective studies. The adopted approach provided evidence for a shift to glucose-based metabolism and high proliferation in adenocarcinoma tissues and identification of chromatin remodelling complexes involved in mouse T-cell activation. The results correlate with the original interpretation of the experiments and also provide novel biological details about protein complexes affected. ComplexBrowser is, to our knowledge, the first tool to automate quantitative protein complex analysis for high-throughput studies, providing insights into protein complex regulation within minutes of analysis.

A fully functional demo version of ComplexBrowser v1.0 is available online via http://computproteomics.bmb.sdu.dk/Apps/ComplexBrowser/

The source code can be downloaded from: https://bitbucket.org/michalakw/complexbrowser

**Highlights:**

- Automated analysis of protein complexes in proteomics experiments
- Quantitative measure of the coordinated changes in protein complex components
- Interactive visualisations for exploratory analysis of proteomics results

**In brief:** ComplexBrowser is capable of identifying protein complexes in datasets obtained from large scale quantitative proteomics experiments. It provides, in the form of the CFC factor, a quantitative measure of the coordinated changes in complex components. This facilitates assessing the overall trends in the processes governed by the identified protein complexes providing a new and complementary way of interpreting proteomics experiments.

## Introduction

Proteomics has become a method of choice for large scale analysis of biological systems. Recent advancements in multidimensional separation methods, improved instrument speed, sensitivity and resolving power allow for the generation of nearly full proteomes in the scope of 35 hours of analysis, providing quantitative information for more than 12 000 gene products and covering 4-6 orders of magnitude [1, 2].

Coverage of nearly complete proteomes raises great research opportunities, but also challenges, especially in the domain of biological interpretation of the results. Currently, this process remains time-consuming and requires considerable expertise.

Commonly used analysis pipelines involve annotating proteins according to their molecular function, cellular component and biological processes, based on information gathered in Gene Ontology (GO) databases [3, 4]. Further GO-term enrichment methods define over-represented annotations in users data, providing a general understanding of the biological processes affected [5].

Pathway analysis is a different approach for determining protein role, concentrated on its specific biochemical activity. Tools such as IPA®, KEGG or Reactome map proteins to molecular pathways [6-8] and visualise processes those gene products are known to be involved in. A clear advantage of this approach is that pathway databases are mostly based on experimental, manually curated data, while the majority of GO-annotations come from *in silico* predictions and text mining [9].

An alternative practice, often employed in studies of less researched organisms, is protein domain and motif analysis [10]. This strategy utilises sequence alignment and secondary structure prediction tools to find similarity between a protein of interest and better-annotated analogues in other species. Identification of a specific sequence motif enables assigning function to previously undescribed proteins.

Analysis of protein-protein interactions (PPI) is a complementary approach used very often in parallel to the above methods. Platforms such as STRING [11] utilise information from co-expression studies, cross-species predictions, experimental evidence and literature mining to build protein interaction graphs, in which nodes represent gene products and edges correspond to interactions. These maps facilitate identification of genes involved in similar processes or influenced by common regulators. Platforms for PPI investigation provide comprehensive information on proteins function and involvement in various biological processes. However, the enormous knowledgebase of these platforms and the amount of interactions drawn from large gene/protein lists is often very difficult to handle and interpret.

Protein complexes are molecular machines that perform many of the key biochemical activities essential to cell e.g. replication, transcription, translation, cell signalling, cell-cycle regulation and oxidative phosphorylation. Their role in maintaining cell homeostasis and involvement in disease development [12] prove that the detailed characterization of protein complex expression would be very helpful in understanding the often highly intertwined processes in the cell.

Interrogation of the expression of components of known protein complexes in large scale proteomics data had been performed in a number of studies [13, 14] https://www.biorxiv.org/content/early/2018/07/11/367227). It is now clear that the majority of known protein complexes are translationally and post-translationally regulated and therefore exhibit co-expression when compared across cell types and tissues. However, so far no automated and user-friendly approach for the analysis of complex behaviour in new datasets had been developed.

In this manuscript we present ComplexBrowser that enables automated and quantitative analysis of protein complexes in proteomics experiments. The software interrogates CORUM [15] and EBI Complex Portal [16] databases to find known protein complexes present in a given protein list and utilises quantitative proteomics data and factor analysis to summarise the overall expression trends for each complex across the studied biological conditions. The re-analysis of two, previously published, large scale proteomics datasets shows the high potential of the approach to gain in-depth knowledge about regulation of protein complexes in different biological contexts.

## Experimental Procedures

### Protein complex databases

The presented software relies on information from two established, manually curated protein complex databases: CORUM [15] and EBI Complex Portal [16] with 2693 and 2454 entries respectively, together covering 22 species (state for 24.05.2018).

### Software design

#### Data Input

ComplexBrowser accepts data tables in .csv or .txt format as input. The file must contain single, unique UniProt [17] accessions in the first column and quantitative information in subsequent columns (LFQ or summarised reporter ion intensities for each analysed sample). Optionally, columns with confidence scores from statistical tests result can be appended. These values are calculated in relation to the first condition appearing in the input file. If they are not included by the user, differential expression analysis using limma [18] is conducted, along with FDR estimation using the qvalue R package [19].

#### Quality control

Prior to the analysis of protein complexes, the software creates data visualisations for quality control (QC) evaluation purposes. The QC visualisations include:

- Boxplot of log-transformed intensities to control for inconsistency in e. g. injection amounts.
- Missing value bar plots generated by summing up the number of missing values in each quantitative column to compare protein coverage between samples.
- Pairwise Scatter plots of log-transformed intensities of all proteins quantified in two selected conditions, displaying sample to sample correlations (Pearson, Kendall, Spearman), to test similarity between the samples.
- Histograms of coefficient of variation (CV) of protein absolute intensity measurements within each experimental condition to assess replicate variation.
- Q-value charts counting the number of differentially expressed features in relation to a set threshold.
- Volcano plots to depict the relation between fold-changes and confidence for differentially regulated features.
- PCA results for visual comparison of all samples.

The software also implements four common, previously described normalisation methods [20].

#### Complex expression analysis

ComplexBrowser employs the FARMS algorithm to determine complex fold change (CFC). FARMS [21] is based on Bayesian factor analysis with an assumption of Gaussian measurement noise. It has previously been used to estimate protein abundance, based on peptide concentrations in the protein summarisation process. It has proven useful in detecting outliers in peptide expression profiles and limiting their influence on protein quantitation [22]. In ComplexBrowser, we employed our own implementation of the FARMS algorithm for performing weighted summarisation of log-transformed expression changes of protein complex subunits. The complex expression is calculated in two steps. First, scaled intensities of all subunits are summed within each replicate, subsequently, the sums are averaged to obtain one value of complex expression for each analysed condition. Relative change of complex expression between two given conditions is defined as complex fold change (CFC). ComplexBrowser also provides a summary measure, describing the amount of variability in the expression profile of a given complex. This information is presented as signal to noise ratio, or in short-noise. Noise of 0 indicates perfect co-expression, while noise value of 1 points to very poor correlation. The default noise threshold set in the software is 0.5.

#### Linearity of complex components co-expression

Building on the idea of using linearity of subunit co-expression as a measure of data quality [23], ComplexBrowser draws supplementary visualisations to investigate co-regulation between different conditions. For a selected protein complex, it takes log-transformed abundances of all its subunits in two conditions and displays them on a scatter plot, where each point corresponds to one protein. Orthogonal distance regression (ODR) is employed to determine the quality of co-expression similarity, since unlike ordinary least squares regression, ODR considers variability in both x and y values, therefore it fits a model that minimizes errors of both measurements [24]. The procedure returns a single R^2^ value per complex for each pair of conditions as a measure of co-expression.

#### Heatmaps and hierarchical clustering

Combined with dendrograms produced by hierarchical clustering algorithms, heatmaps allow detecting proteins not following the common trend of a protein complex. These could be subunits that participate in several different complexes or interact with a given complex transiently. ComplexBrowser displays two different heatmaps – expression and correlation (between protein expressions). The expression heatmap displays log-transformed, mean-normalised expression values for all subunits within selected complex across all experimental conditions. The same data input is used to calculate pairwise correlations between expression profiles of all complex subunits. Based on this infromation a correlation matrix is computed and displayed as the correlation heatmap. Both graphs use aggregative hierarchical clustering implemented in R’s hclust function to provide dendrograms. The function uses distance measures and linkage functions selected by the user in the graphical interface.

### Test datasets

ComplexBrowser performance was tested using protein quantification data from two previously published studies [25, 26]. Further on in the manuscript we refer to the studies as *adenocarcinoma dataset* [26] and *T-cell dataset* [25].

*Adenocarcinoma dataset* investigates protein expression differences between formalin-fixed, paraffin embedded tissue samples from patients with colon cancer in comparison to healthy colon mucosa and nodal metastatic tumours using label-free quantitation based on LFQ intensities. The MS proteomics data of the *adenocarcinoma dataset* were obtained from supplementary tables of the original publication available from the publisher’s site [26]. For the purpose of this analysis we have discarded samples denoted as “CA2” and “NO2” to ensure an equal number of replicates in each condition. We have filtered the protein intensities table to retain proteins with at least 4 valid quantitative values within each condition. We have removed isoform identifiers from the original accession numbers and rows with non-unique identifiers were removed. This resulted in a dataset containing LFQ values for 6824 proteins from 3 conditions, 7 biological replicates each. The input file used is this study can be found in supplementary Table S1.

The *T-cell dataset* studies activation of quiescent mouse T cells over four time points (0, 2, 8 and 16h) in two biological replicates. Proteins were quantified using tandem mass tags (TMT) labelling and were analysed on an Orbitrap Elite MS instrument. The data were obtained from the original publication from PRIDE database (accession numbers PXD004367 and PXD005492) [27]. The dataset contained normalised intensities for 8431 proteins. The input file used is this study can be found in Table S2.

### Software implementation

ComplexBrowser was implemented in R [28]. The user interface was developed using Shiny, Plotly, networkD3, heatmaply, DT and data.table libraries, allowing interactive and adjustable data visualization. PreprocessCore, stringr, pracma, dplyr, limma and qvalue packages were used for data manipulation and statistical analysis.

### Software accessibility

The tool can be accessed via web service at http://computproteomics.bmb.sdu.dk/Apps/ComplexBrowser or can be run locally after installation of Rstudio and required libraries. For detailed instructions see supplementary File S1.

The source code can be downloaded from: https://bitbucket.org/michalakw/complexbrowser

## Results

We have developed ComplexBrowser for analysis of protein complexes abundance and co-expression in large scale proteomics experiments. The general analysis pipeline implemented in the program is presented in Fig. 1. An extensive description of ComplexBrowser’s procedures and results can be found in supplementary File S1. In brief, a table containing the quantitative information of the identified proteins is uploaded using the web browser interface. After defining parameters of the analysis (e.g. number of conditions and replicates) the analysis of the quality of the quantitative data provided is carried out and visualised. In a following window analysis of p the presence and changes in abundance of protein complexes is carried out. Interactive tables and graphics allow the user to conveniently evaluate the results of the analysis. Tables containing results and a summary report are available for download.

**Figure 1.**
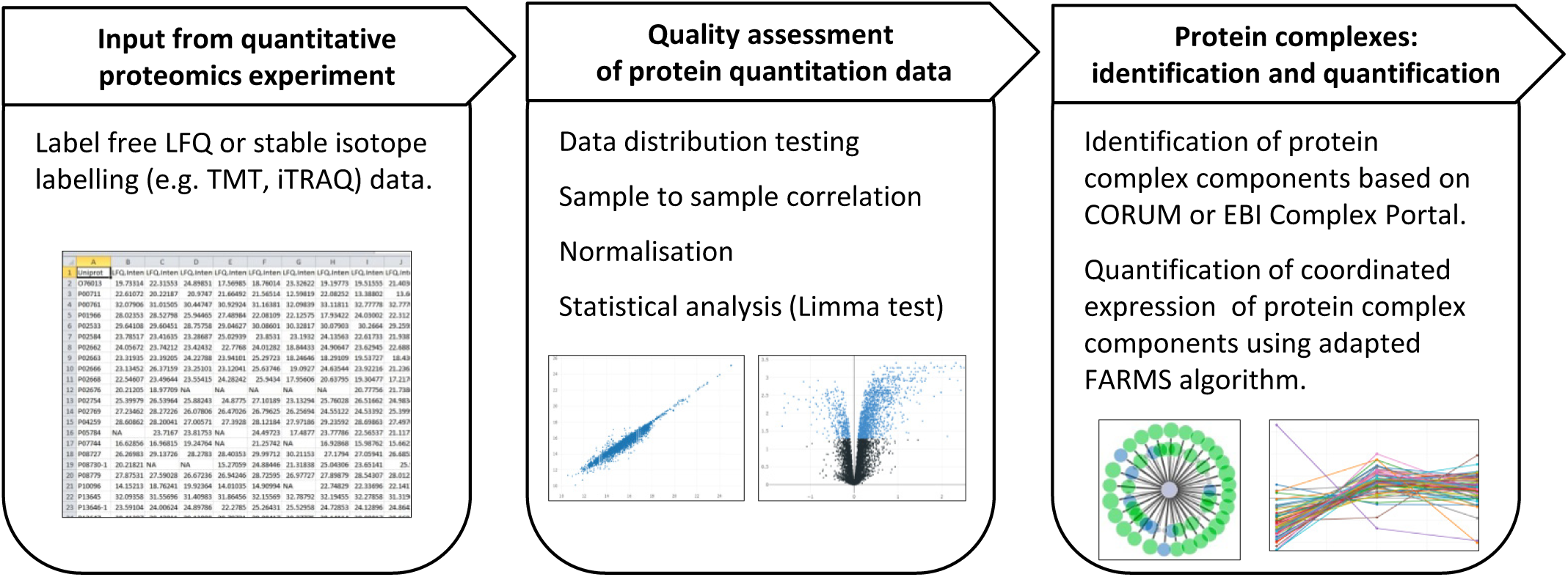
ComplexBrowser analysis workflow. The list of identified proteins along with quantitative information is loaded to the software and analysis parameters are set. This is followed by the assessment of the quality of the quantitative data provided. Finally presence and changes in abundance of protein complexes is interrogated.

To test the performance of the developed platform, we have used two published proteomics studies: *adenocarcinoma dataset* [26] and *T-cell datas*et [25]. The results of quality control and protein complex analysis steps are presented below.

### Quality control of proteomics data in ComplexBrowser

*Adenocarcinoma dataset* containing quantitative proteomics values from 3 biological conditions with 7 replicates each were uploaded to ComplexBrowser in the following order: C1 – control, C2 – metastasis, C3 – cancer. A summary of the quality analysis step can be found in a supplementary File S2.

Analysis of Boxplot graphs of log-transformed intensities, Fig. 2A, indicated that normalisation was necessary to reduce the variability between intensity distributions and ensure sample to sample comparability; therefore a quantile normalisation was performed. Despite the normalisation, the mean CV values were 65, 78 and 77% for normal, metastasis and cancer samples respectively, Fig. 2B, indicating relatively large variability within measurements, most likely due to the clinical character of the samples.

**Figure 2.**
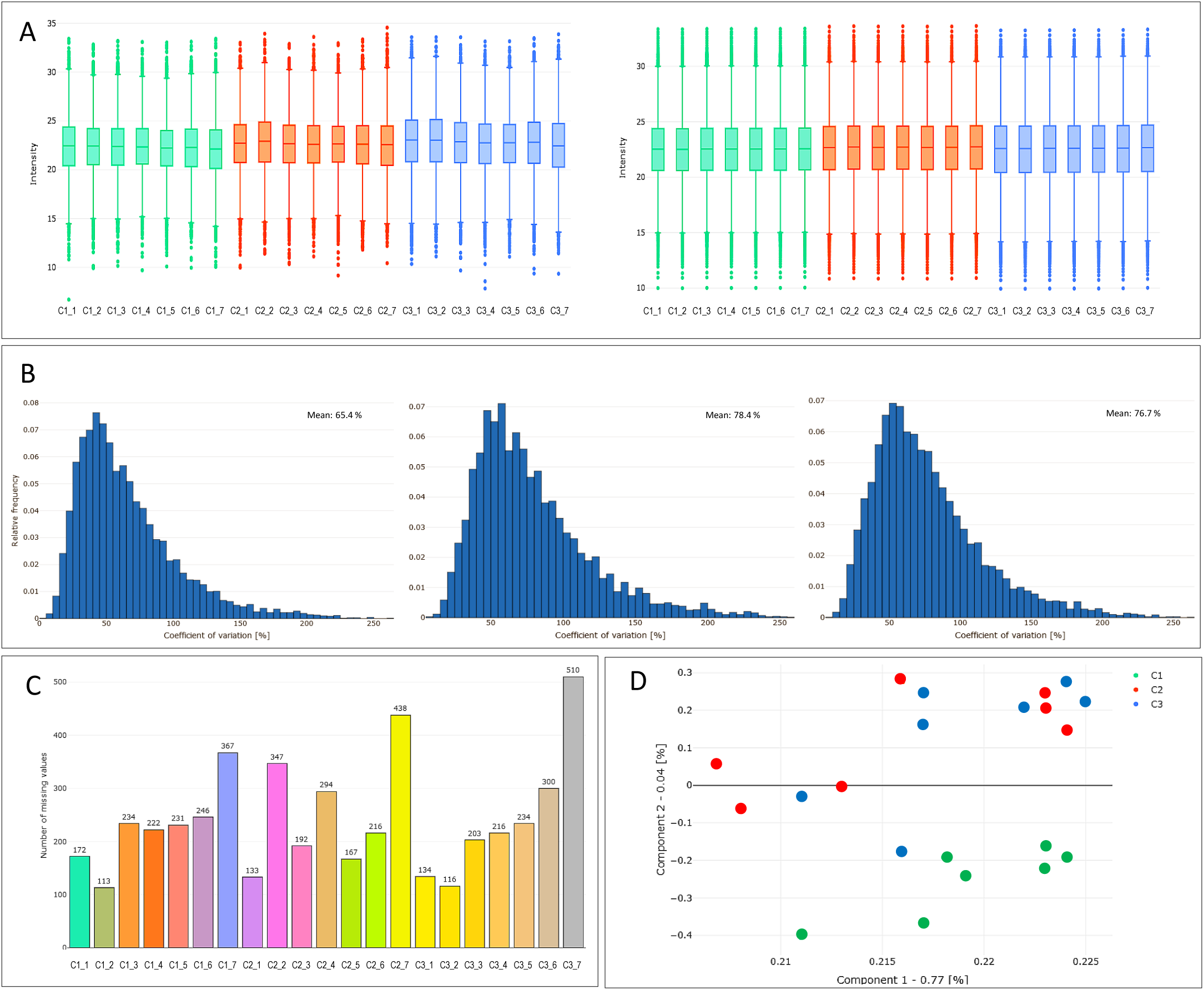
Examples of data quality analysis visualisation of the Adenocarcinoma dataset using ComplexBrowser. A - Box plot of log-transformed LFQ intensities of all identified proteins in each sample pre- (left panel) and post- (right panel) normalisation; B - CV distribution of LFQ intensities of all identified proteins within each analysed condition; C – Column graph representing the number of missing values observed in each sample; D - Principal component analysis based on all identified and quantified proteins. Component 1 and 2 with [%] of variance explained; C1 –normal, C2 – cancer and C3 – metastatic tissue; C1_1, C1_2 etc. depict different samples with in each condition.

Label free experiments may contain proteins that have been quantified only in few samples and in other samples their quantitative value is missing (referred to as “missing values”). Many missing values in few samples could indicate lack of technical reproducibility and/or quality of the obtained data. In the *adenocarcinoma dataset* the number of missing values per sample varied from 113 to 510 and consisted in total only 3.5% of all valid measurements, Fig. 2C. It did not show any persistent bias of the data.

Differentially expressed features were determined using a paired limma test and FDR estimation, suplementary Table S3. Taking into account large deviation in measurements and clinical sample character, features with 0.01 FDR value were considered, resulting in identification of 802 proteins differentially regulated for metastasis and 813 for cancer samples. PCA analysis has shown a good separation of control samples but an overlap between the carcinoma and metastatic tissues, Fig. 2D.

Subsequently ComplexBrowser data quality analysis was applied to the *T-cell dataset*, which consists of four sets (0, 2, 8 and 16 h) of two replicates and was generated using TMT-based MS^2^ quantitation. The complete report can be found in supplementary File S3. Boxplots graphs distributions of *T-cell dataset* results had shown only very small variation in median values for the different samples and did not require any further normalisation, supplementary File S3. No missing values were observed. The variability of measurements was noticeably lower in comparison to *Adenocarcinoma* experiment with average CV from all conditions of 4.56% (vs 73.51% for the previous dataset), supplementary File S2 and S3. Fig. 3A illustrates high level of sample to sample correlation of TMT intensities for each protein between selected samples. An increasing number of differentially expressed proteins (39, 1869 and 5600) were detected after 2, 4 and 16 h of T-cell activation, at FDR of 0.05. Fig.3 3B presents volcano plots generated by the ComplexBrowser software. Results of statistical analysis can be found in supplementary Table S4.

**Figure 3.**
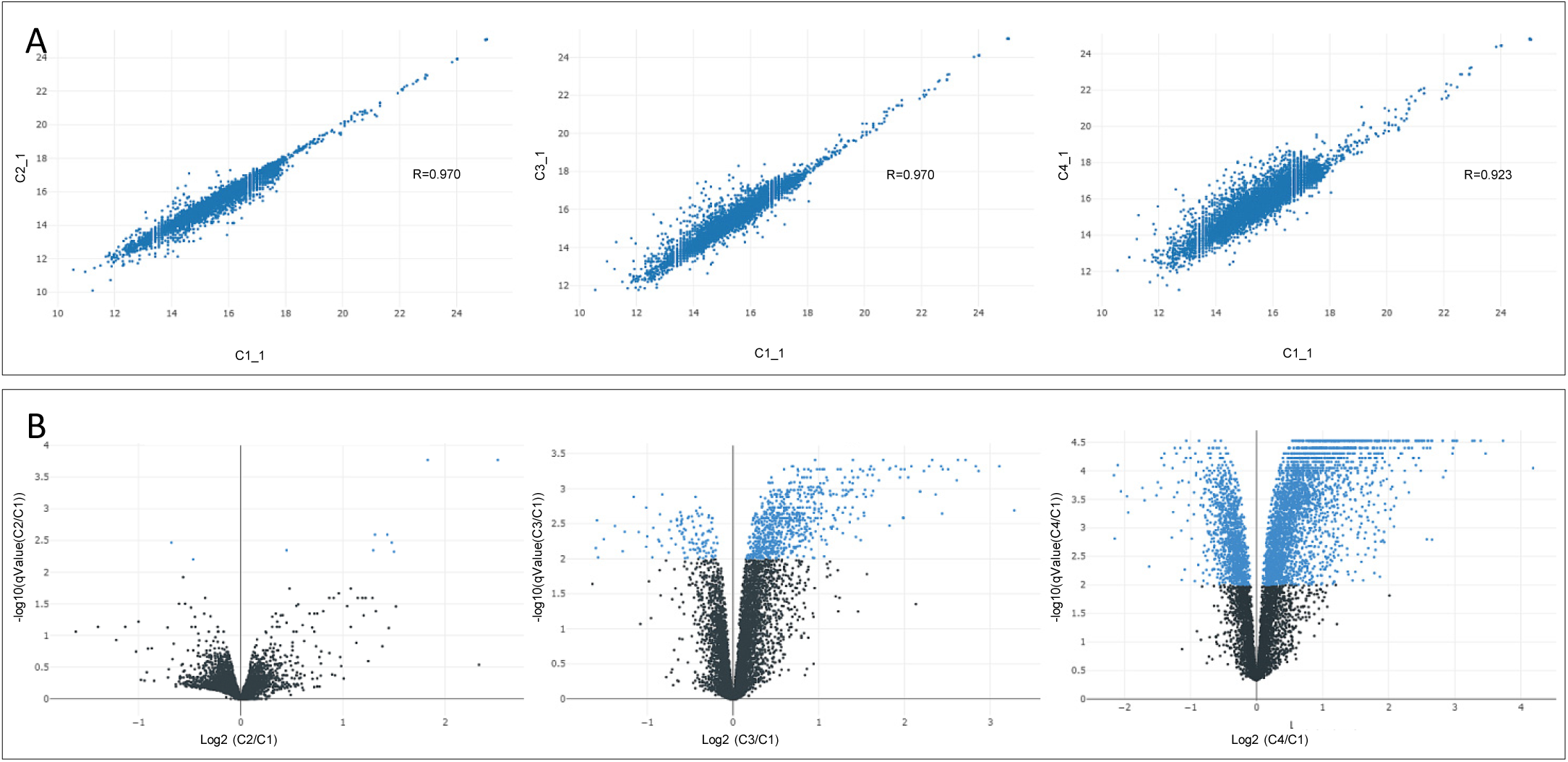
Example of ComplexBrowser generated visualisation of quality control analysis of the T-cell dataset. A – Pearson correlation of TMT intensities between 4 selected samples. Left panel - C1_1 vs. C2_1; Middle panel - C1_1 vs. C3_1; Right panel - C1_1 vs. C4_1; B – Volcano plots at FDR of 0.05; C1, C2, C3 and C4 depict non stimulated T- cells (0 h) and T-cells stimulated for 2 h, 8 h and 16 h respectively; C1_1 and C1_2 etc. depict different replicates within each condition.

### Protein complex analysis

ComplexBrowser facilitates the analysis of protein complex expression in large-scale studies. The software queries the input data for proteins reported to be complex members, investigates their co-expression patterns and visualises results according to specified statistical, noise and expression change thresholds. It quantifies changes of complex abundance by calculating the complex fold change factor (CFC). Resulting graphs and tables allow for data exploration using interactive figures.

We recommend that setting parameters for complex analysis should be performed based on the results obtained from the quality analysis module. The *adenocarcinoma dataset* showed a relatively high variation between samples therefore it was analysed using the CFC≥1.5, complex Noise ≥ 0.5 and protein expression q value ≤ 0.05. The *T-cell dataset* had shown lower measurement variability and good correlation between the biological replicates therefore the CFC was set to ≥1.2, complex Noise threshold was kept at ≥ 0.5 and protein expression q value ≤ 0.05.

The main results of the complex analysis module are represented in form of a tabular output, supplementary Table S5 and S6, using CORUM and EBI databases respectively), which can be downloaded for use in external applications. In ComplexBrowser the table can be sorted to provide easy access to the relevant complexes. It contains complex ID, complex names, number of proteins (subunits) of the complex identified and quantified in the analysed dataset, number of all subunits and a % complex coverage that allows the user to identify complexes that are highly represented in the analysed dataset. CFC – complex fold change, together with Noise factor, is given for the analysed conditions providing means to evaluate the coordinated changes in expression of complex components. Accession numbers of all identified complex subunits, gene ontology annotations for the complex are also listed in the table.

#### Protein complexes identified in *adenocarcinoma* dataset reflect biological features of cancer and metastatic tissues

Protein complex analysis of *adenocarcinoma* dataset identified 1519 protein complexes from CORUM and 366 from Complex Portal Table 1. Typical visualisation of a protein complex and its components generated by ComplexBrowser is presented in Fig. 4. The top 5 most upregulated and top 5 most downregulated complexes in cancer tissue selected based on the CFC are shown in Table 2. Mitochondrial respiratory chain I (−4.368 CFC), F1F0-cytochrome C oxidase (−4.188 CFC) were identified as the key complexes downregulated in both metastatic and cancer samples, Table 2. This finding combined with a significant down regulation of ATP synthase (−1.263 CFC) indicates a shift in cancer cells metabolism from oxidative phosphorylation to glycolytic pathways, which is consistent with the results of the original publication [26].

**Table 1.**
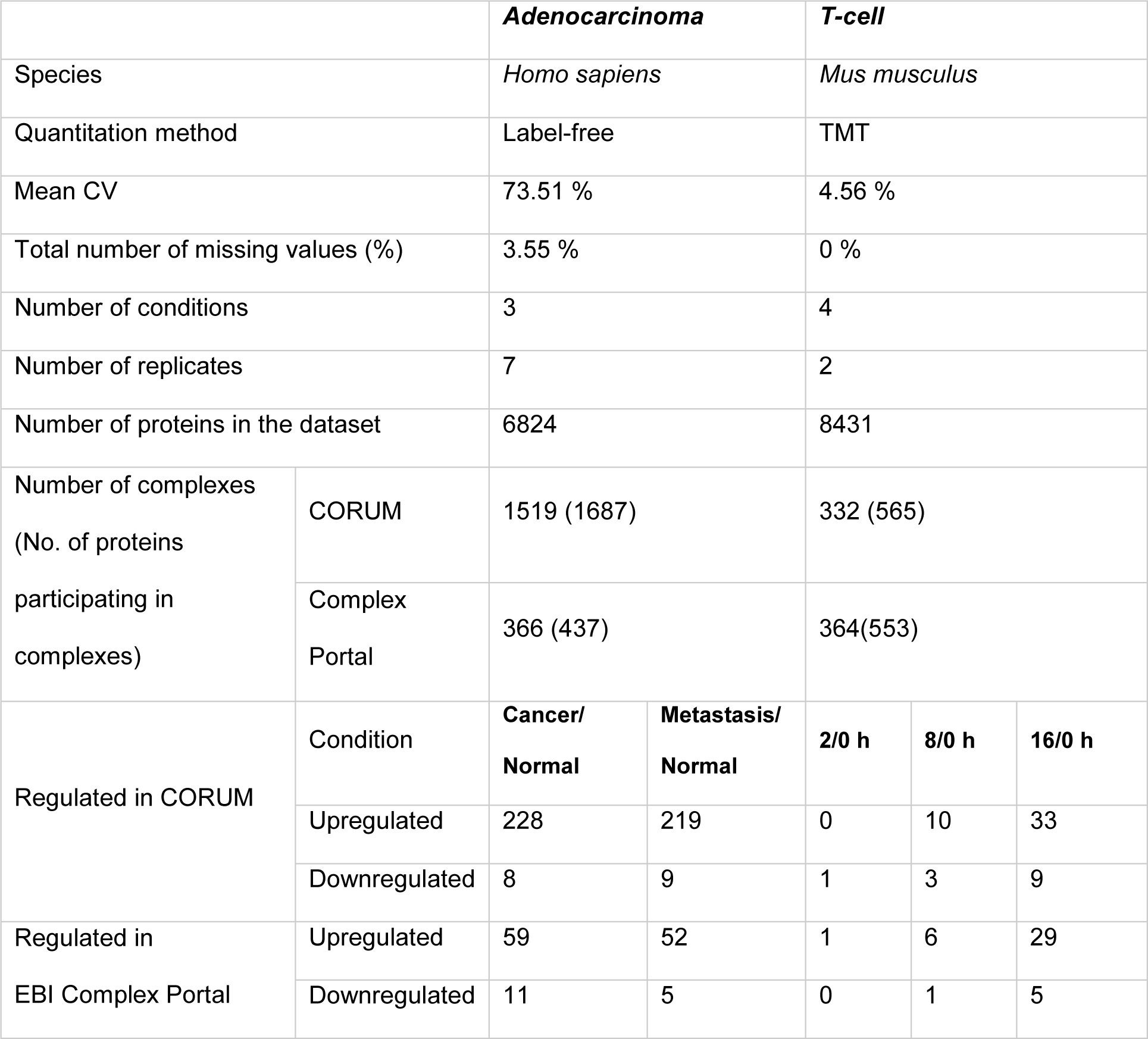
Summary of ComplexBrowser based analysis of protein complexes in Adenocarcinoma (CFC≥1.5, complex Noise ≥ 0.5, protein expression q value ≤ 0.05) and T-cell datasets (CFC was set to ≥1.2, complex Noise ≥ 0.5, protein expression q value ≤ 0.05).

|  |  | **Adenocarcinoma** | **T-cell** |  |  |  |
| --- | --- | --- | --- | --- | --- | --- |
| Species |  | *Homo sapiens* | *Mus musculus* |  |  |  |
| Quantitation method |  | Label-free | TMT |  |  |  |
| Mean CV |  | 73.51 % | 4.56 % |  |  |  |
| Total number of missing values (%) |  | 3.55 % | 0 % |  |  |  |
| Number of conditions |  | 3 | 4 |  |  |  |
| Number of replicates |  | 7 | 2 |  |  |  |
| Number of proteins in the dataset |  | 6824 | 8431 |  |  |  |
| Number of complexes<br>(No. of proteins<br>participating in<br>complexes) | CORUM | 1519 (1687) | 332 (565) |  |  |  |
|  | Complex<br>Portal | 366 (437) | 364(553) |  |  |  |
| Regulated in CORUM | Condition | **Cancer/**<br>**Normal** | **Metastasis/**<br>**Normal** | **2/0 h** | **8/0 h** | **16/0 h** |
|  | Upregulated | 228 | 219 | 0 | 10 | 33 |
|  | Downregulated | 8 | 9 | 1 | 3 | 9 |
| Regulated in<br>EBI Complex Portal | Upregulated | 59 | 52 | 1 | 6 | 29 |
|  | Downregulated | 11 | 5 | 0 | 1 | 5 |

**Table 2.**
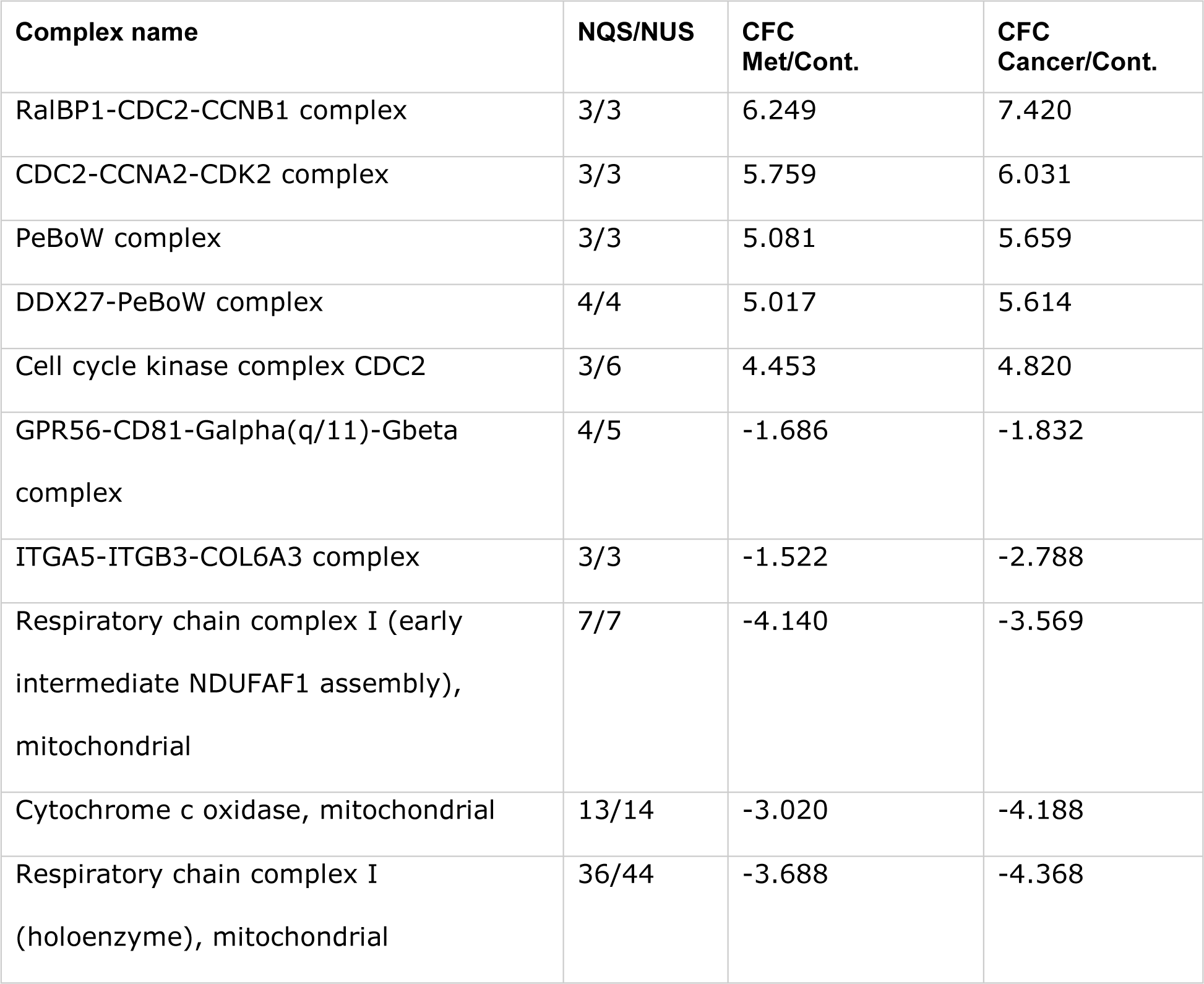
Top 5 up and downregulated protein complexes in Adenocarcinoma dataset based on CORUM database; NQS - number of quantified subunits; NUS - number of unique subunits; CFC Met/Cont. – complex fold change between metastatic and control samples; CFC Cancer/Cont. - complex fold change between cancer and control samples;

**Figure 4.**
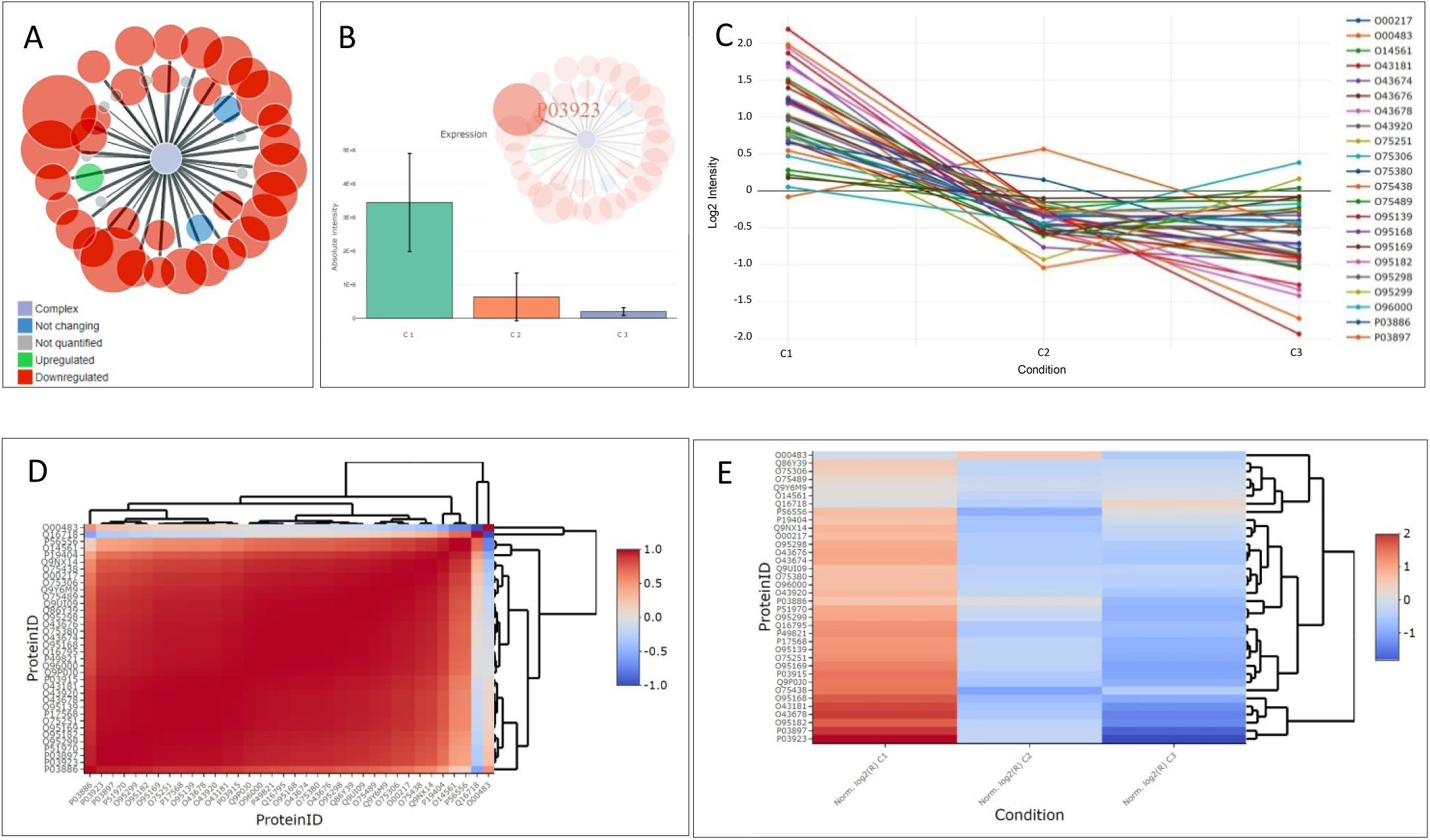
Visualisation of Respiratory Chain Complex I (holoenzyme) identified in Adenocarcinoma dataset using ComplexBrowser; A - Star graph representing complex components and their changes between metastasis and control samples; nodes (circles) denote subunits of the complex; the size of the nodes represents the degree of fold change; the colours indicate the type of regulation (red – downregulated, green – upregulated, blue – not changing, grey – not identified); thick edges (lines) indicate differentially regulated proteins; B - Expression of NADH-ubiquinone oxidoreductase chain 6 (SwissProt: P03923) a subunit of Respiratory Chain Complex I showing decreased expression in cancer and metastasis tissues; C - Expression profiles of all identified and quantified subunits of the Respiratory Chain Complex I presented in panel A; D – Pearson correlation heat map visualising log-transformed, mean normalised intensities of all identified and quantified subunits of the Respiratory Chain Complex I presented in panel A in the three analysed conditions. E – Heat map visualising log-transformed, mean normalised intensities of all identified and quantified subunits of the Respiratory Chain Complex I presented in panel A in the three analysed conditions; C1 – normal, C2 – cancer and C3 – metastatic tissue;

Among the top 5 regulated complexes we have identified overexpression of 3 complexes related to mitotic cell cycle progression and activation: RalBP1-CDC2-CCNB1 complex (7.420 CFC), CDC2-CCNA2-CDK2 complex (6.031 CFC); and Cell cycle kinase complex CDC2 (4.820 CFC), for more details see Table 2 and supplementary Table S4. This points to an increased activation of cell division and replication, as well as reduction in respiration, which are known characteristics of cancer [29, 30].

In addition to previously described results, ComplexBrowser detected 3.58 fold upregulation of MTA1 complex involved in metastatic tumour formation in nodal tissue samples [31]. The same complex was changed 2.33 fold in main tumour tissue supplementary Table S4.

#### Activation of mouse T-cells is reflected in coordinated changes of protein complex components

ComplexBrowser was used to analyse protein complexes during activation of murine T-cells. The dataset consists of 4 consecutive time points (0, 2, 8 and 16h) collected after activation. It is expected that trends of coordinated changes of protein complexes would follow the timeline of the activation events and would be reflected in the CFC changes.

Despite the large number (8431) of proteins present in the *T-cell dataset* complex analysis had identified only 332 and 374 protein complexes in the CORUM and EBI Complex Portal, Table 2. This is most likely because there is fewer mouse specific protein complexes present in those databases. Additionally the complexes identified with the two databases are very different and share only 16% of the involved proteins.

Despite those drawbacks use of both databases identified an increasing number of significantly changing complexes was over the different time points of T-cell activation reflecting the gradual changes in the proteome of activated cells. For example, based on EBI Complex Portal, one complex was significantly regulated 2 hours after T-cell activation and 7 and 34 at 8 and 16 h time points respectively, Table 2. A summary of top 5 most upregulated and top 5 most downregulated complexes after 16 h of T-cell activation can be found in Table 3.

**Table 3.**
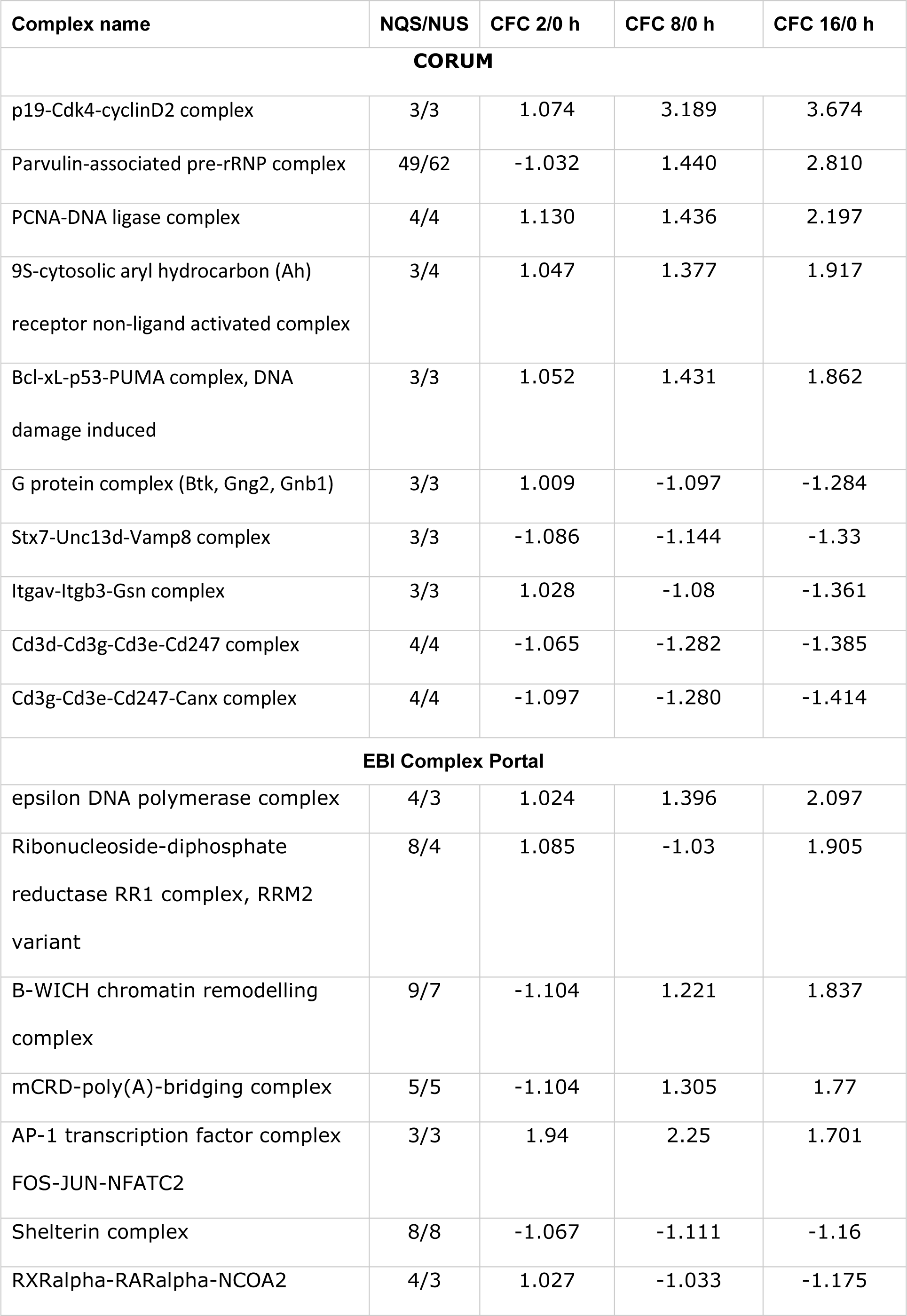
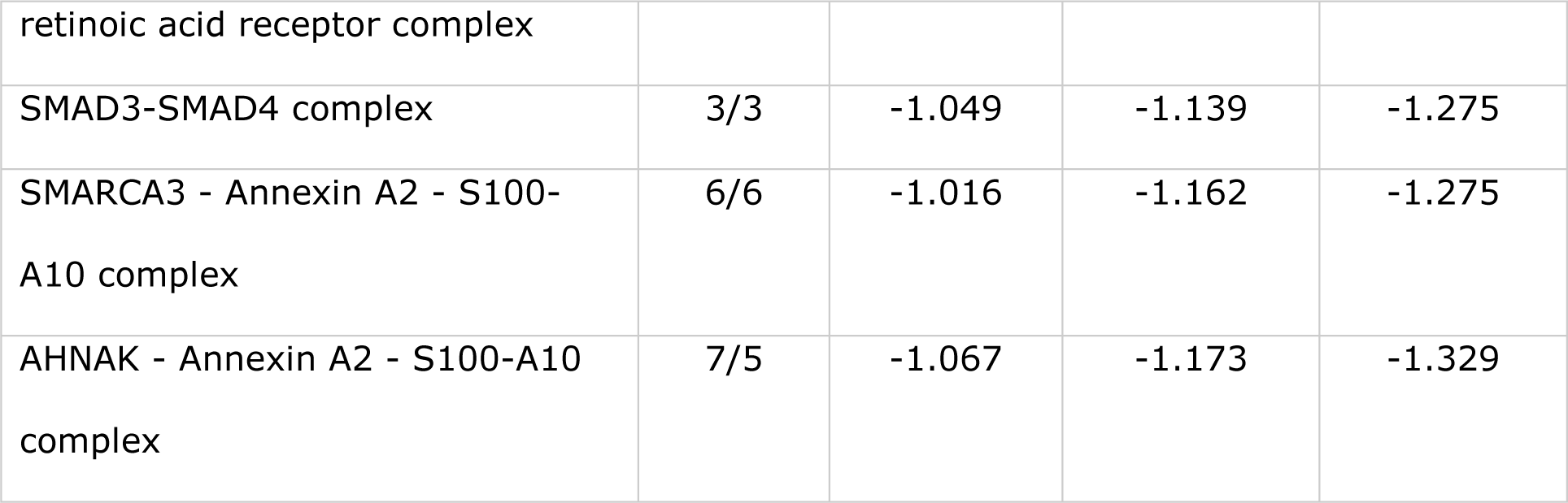
Top 10 up and downregulated protein complexes in T-cell experiment 16 h after activation based on CORUM and EBI Complex Portal databases, result for 2 h and 8 h after activation are shown for comparison; NQS - number of quantified subunits; NUS - number of unique subunits; CFC 2/0 h, CFC 8/0 h and CFC 16/0 h – complex fold change between 2, 8 and 16 h after activation and control samples (0 h) respectively.

The largest quantified complex was Parvulin-associated pre-rRNP complex (CORUM) with 49 out of 62 subunits quantified, Fig. 5A. Its abundance was highly correlated (Noise = 0.001) and increased over the time course of the experiment (−1.03, 1.44, 2.81 CFC after 2, 8 and 16h of activation respectively). This protein complex is involved in ribosome biogenesis and contains several ribosomal subunits [32]. BAT3 complex (EBI Complex Portal), responsible for the targeting of the transmembrane domain-containing proteins from the ribosome towards membranes, was also upregulated in the last condition (CFC 1.42), Fig. 5B, together with pre-ribosomal proteins overexpression suggesting an increase in protein synthesis.

**Figure 5.**
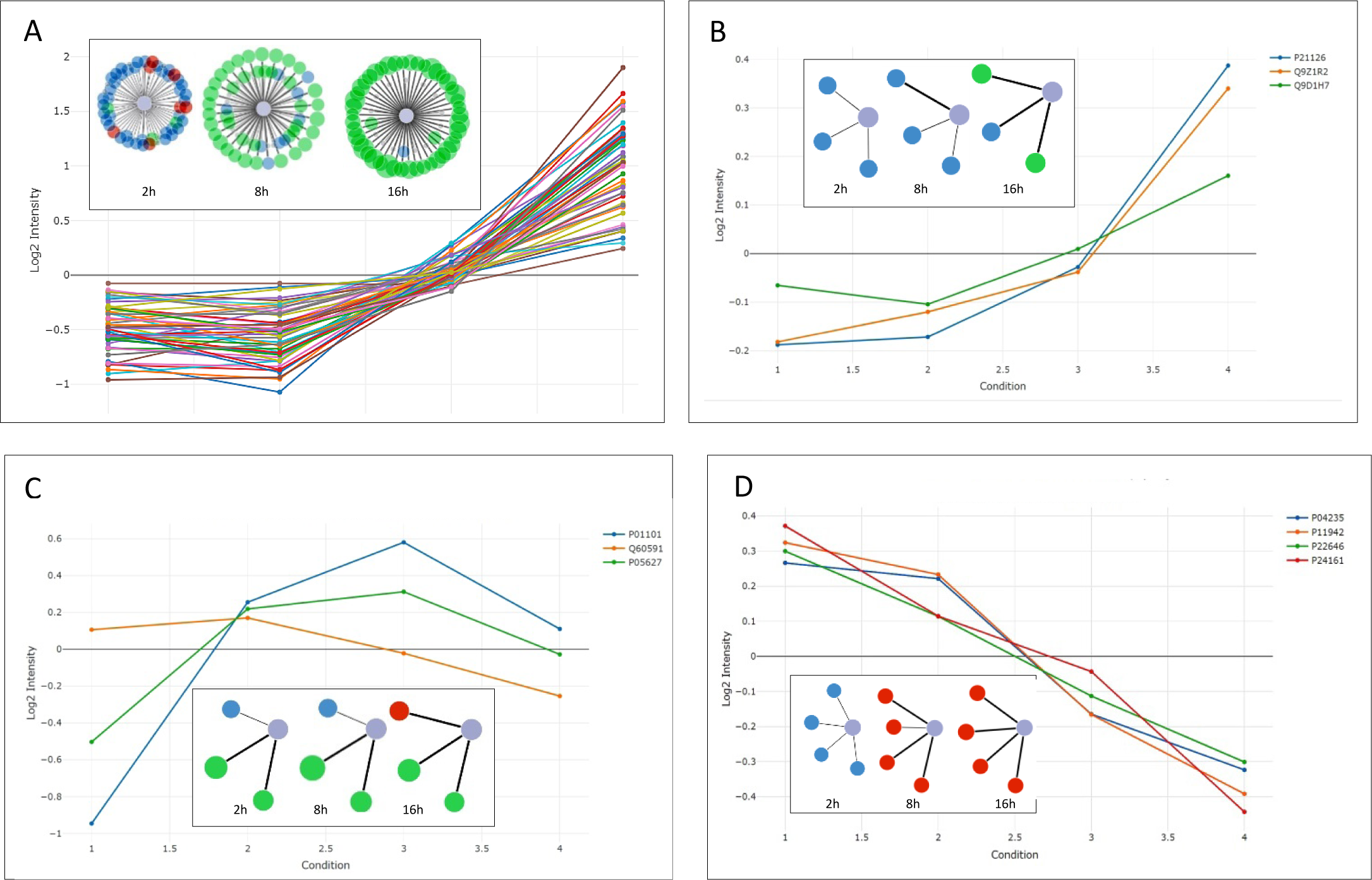
Visualisation of selected protein complexes identified in T-cell dataset using ComplexBrowser. Inserts contain star graphs representing complex components and their changes during T-cell differentiation; nodes (circles) denote subunits of the complex; the size of the nodes represents the degree of fold change; the colours indicate the type of regulation (red – downregulated, green – upregulated, blue – not changing, grey – not identified); thick edges (lines) indicate differentially regulated proteins. Graphs are normalised expression profiles visualising the expression of the subunits of complexes at 2, 8 and 16 hours after T-cell activation. C1, C2, C3 and C4 depict non stimulated T-cells (0 h) and T-cells stimulated for 2 h, 8 h and 16 h respectively. A - Parvulin-associated pre-rRNP complex; B - AP-1 transcription factor complex FOS-JUN-NFATC2; C – BAT3 complex; C – Cd3d-Cd3g-Cd3e-Cd247 complex

Combined data from CORUM and EBI Complex Portal showed that 62 complexes were induced 16 h after activation, with a substantial increase in abundance between 8 h and 16 h after activation, supplementary Table S7 and S8.

p19-Cdk4-cyclinD2 (3.96 CFC after 16 h) and Cyclin D1-associated (2.47 CFC after 16 h) protein complexes were found amongst the most upregulated after 8 and 16 hours after activation. These two complexes are involved in the regulation of cell cycle and transition through G1 phase and their coordinated upregulation reflects the transition of T-cells into proliferative state. These changes were accompanied by increase in the expression of various DNA polymerase complexes e.g. DNA synthesome (1.49 CFC after 16 h) and DNA polymerase alpha, delta and epsilon complexes (1.45, 1.25, 2.09 CFC after 16 h) and Telomerase holoenzyme complex (1.40 CFC after 16 h) pointing to increase in processes associated with DNA replication and cell proliferation.

AP-1 transcription factor complex FOS-JUN-NFATC2 (CORUM) (CFC 1.87 at 2h, 2.13 at 8 h and 1.62 at 16h) is an example of a complex with highly coordinated early response to the stimulus, Fig. 5C. Similar patterns in the AP-1 complex were also reported by Tan *et al.* [25].

Down regulation trend was found in the expression of the Cd3d-Cd3g-Cd3e-Cd247 complex (1.07, −1.2, −1.36 CFC), a membrane glycoprotein assembly which is a part of the T-cell co-receptor, Fig. 5D, that is shut down during T-cell activation [25].

Additionally we have identified a variety of complexes involved in post-translational modification of histones and chromatin remodelling majority of which showed increased expression at 8 and 16 h of T-cell activation, Table 4. These complexes and regulation of epigenetic processes in maturation of T-cell had been largely overlooked in the original study. Describing the regulation of transcription by post-translational modifications and chromatin remodelling requires their quantification which was not done in this case; therefore an extensive follow-up analysis might be necessary to interpret these results.

**Table 4.**
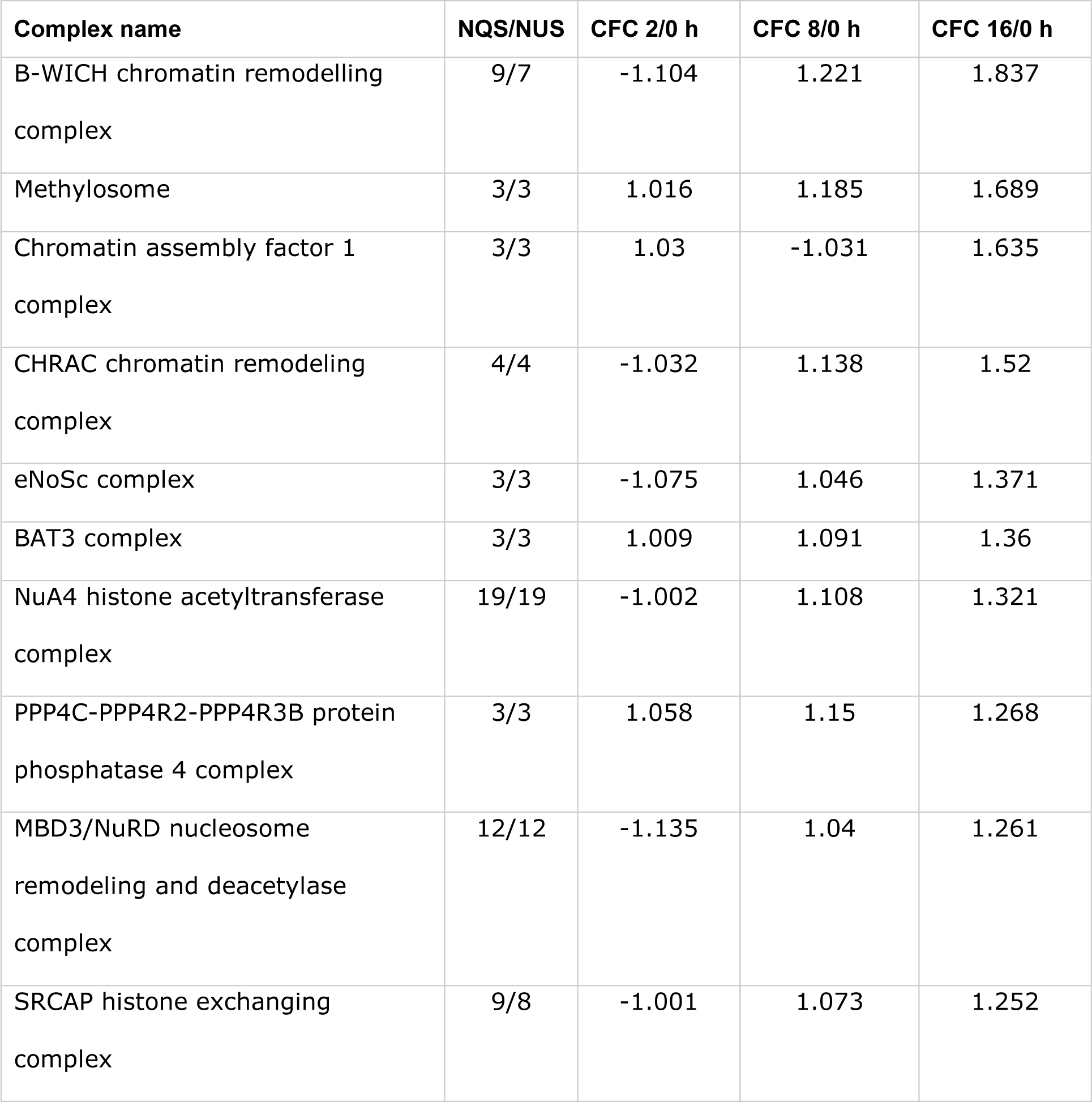
Chromatin remodelling and histone modifications associated protein complexes identified in T-cells 16 h after activation; NQS - number of quantified subunits; NUS - number of unique subunits; CFC 2/0 h, CFC 8/0 h and CFC 16/0 h – complex fold change between 2, 8 and 16 h after activation and control samples (0 h) respectively.

In T-cell dataset ComplexBrowser analysis identified upregulated complexes involved in DNA replication and chromatin remodelling, protein synthesis initiation and cell cycle progression, which allowed us to conclude that the T-cells undergo significant cellular reprogramming and exit a quiescent state 8 to 16 hour after stimulation. Moreover, there was a decrease in T-cell signalling receptors expression from the first time point of the experiment.

## Discussion

ComplexBrowser is, to our knowledge, the first automated tool that enables quantitative analysis of protein complexes in proteomics experiments. It is available through a web-browser and does not require any installations or programming experience. Thus, it has high potential for integration into data analysis workflows commonly used by the scientific community.

Based on the datasets analysed in this manuscript we have shown that it is compatible with label free and TMT based quantitation experiments. And, there are no technical limitations that prevent it from being used with any other MS based protein quantitation method or even gene expression data. ComplexBrowser can handle large proteomic studies with over 8000 quantified proteins and is able to display summary results within one minute from data input. Interactive visualisations provide an intuitive tool for exploratory analysis and data interpretation, enabling the user to investigate the behaviour of whole complexes, as well as single subunits.

The CFC effectively assists in finding complexes that are changing expression in a synchronised manner and is a measure of complex behaviour. Subunits which are not coherent with trends in complex expression are also easily identified using the extensive visualisation tools implemented in the software.

In both test datasets ComplexBrowser identified the key protein complexes that are known to be regulated in cancer and during T-cell activation. The biological interpretation enabled by interrogating of protein complexes was in agreement with the conclusions drawn based on GO-annotations and existing literature. The tool added new insights based on the investigation of annotated protein complexes that were previously not considered in the analysis. Thus ComplexBrowser provides a complementary approach to for example STRING or GO-term enrichment tools. More studies, including different biological perturbations need to be analysed in ComplexBrowser to obtain a complete picture of its utility.

ComplexBrowser relies on information stored in CORUM [15] and Complex Portal databases [16] and is therefore dependent on the efforts of their administrators. The composition of these resources introduces a bias in the analysis, since the largest proportion of complexes described in both databases are of human origin (66.36% of CORUM and 25.79% of Complex Portal). Thus, currently, ComplexBrowser is most suitable for analysis of human proteins. This is visible when comparing the number of proteins found to be involved in complexes from *adenocarcinoma* (human) and *T-cell* (mouse) dataset, Table 1. Aditionally the databases contain entries that are not fully annotated. Further developments of the databases will improve the results provided by the software.

## Supporting information

File S1_ComplexBrowser_manual

File S2_QCreport_Adenocarcinoma

File S3_QCreport_T-cell

## Abbreviations

CFC: Complex fold change
CV: Coefficient of variation
FARMS: Factor Analysis for Robust Microarray Summarization
FC: Fold change
FDR: False discovery rate
GO: Gene ontology
KEGG: Kyoto Encyclopedia of Genes and Genomes
LFQ: Label-free quantitation
LTQ: Linear trap quadrupole
ODR: Orthogonal distance regression
PCA: Principal component analysis
PPI: Protein-protein interaction
STRING: Search Tool for the Retrieval of Interacting Genes/Proteins
TMT: Tandem Mass Tag
QC: Quality control

## Acknowledgments

ARW was supported by a grant from the Independent Research Fund Denmark – Natural Sciences and VILLUM Foundation for a grant to the VILLUM Center for Bioanalytical Sciences at SDU. VS was supported by ELIXIR DK. WM was supported by student grant from MC2 Therapeutics ApS. We thank Ole N. Jensen for his critical comments to the project and manuscript.

## Data Availability

ComplexBrowser’s source code is publicly available at: https://bitbucket.org/michalakw/complexbrowser.

The online version of the application can be found at http://computproteomics.bmb.sdu.dk/Apps/ComplexBrowser/

