## Supplementary material for "ComplexBrowser: a tool for identification and quantification of protein complexes in large scale proteomics datasets": File S1_ComplexBrowser_manual

### ComplexBrowser manual v1.0

ComplexBrowser is a R based software for supervised analysis of changes in protein complex abundance and subunit co-expression in proteomic datasets. It uses information contained in CORUM and EBI Complex Portal databases to provide the user with visualizations helping with biological interpretation of obtained results. ComplexBrowser also incorporates a normalization and quality control steps.

#### Expression change definitions

Due to various terms used differently among published work. Firstly we define following terms for absolute intensities of protein measurement in 2 conditions.

$$I_1 - Absolute\ intensity\ for\ Protein\ X\ in\ codition\ 1\ (CONTROL)$$

$$I_2 - Absolute\ inteisty\ for\ Protein\ X\ in\ condition\ 2\ (TREATMENT)$$

$$Ratio : R = \frac{I_2}{I_1}$$

$$Fold\ change: FC = \begin{cases} R\ for\ R \geq 1 \\ -\frac{1}{R}\ for\ R < 1 \end{cases}$$

$$Log(Ratio): R_L = \log_2\left(\frac{I_1}{I_2}\right) = \log_2(R)$$

#### Data input

ComplexBrowser is accepts data tables in .csv or .txt format as input. The file must containing single, unique UniProt accessions in the first column and quantitative data in subsequent columns. Optionally columns with statistical test result can be added. These values should be calculated in relation to the first condition appearing in the input file. If not included by the user, differential expression analysis using limma(1) will be conducted and FDR(2) estimation using qvalue R package will be provided.

| | ProteinID | 0h.a | 2h.a | 8h.a | 16h.a | 0h.b | 2h.b | 8h.b | 16h.b |
|---|---|---|---|---|---|---|---|---|---|
| A | Q8BGE4 | 23000 | 42000 | 40000 | 32000 | 22000 | 40000 | 35000 | 31000 |
| | Q04211 | 18000 | 51000 | 20000 | 18000 | 19000 | 49000 | 22000 | 18000 |
| | P15037 | 20000 | 46000 | 66000 | 25000 | 19000 | 43000 | 55000 | 28000 |
| | O89113 | 5100 | 8600 | 9100 | 7200 | 5700 | 9200 | 7600 | 7000 |

| | ProteinID | 0h.a | 0h.b | 2h.a | 2h.b | 8h.a | 8h.b | 16h.a | 16h.b |
|---|---|---|---|---|---|---|---|---|---|
| B | Q8BGE4 | 23000 | 22000 | 42000 | 40000 | 40000 | 35000 | 32000 | 31000 |
| | Q04211 | 18000 | 19000 | 51000 | 49000 | 20000 | 22000 | 18000 | 18000 |
| | P15037 | 20000 | 19000 | 46000 | 43000 | 66000 | 55000 | 25000 | 28000 |
| | O89113 | 5100 | 5700 | 8600 | 9200 | 9100 | 7600 | 7200 | 7000 |

Table 1. A Example input with ungrouped replicates columns. B. Example input with grouped replicate columns.

After loading the data user must specify following input parameters:

- Separator used in the file
- Decimal symbol used in the file
- Number of conditions
- Number of replicates (must be equal in all conditions)
- Numerical values format (absolute or log2-transformed)
- Order of quantitative columns (Grouped or not grouped)
- Presence of statistical test results (if not provided, specify paired or unpaired experimental design)
- **Press "Run QC"**

#### Quality control

Due to irreproducibility in sample handling, human error, fluctuation in injection amount and technical variability quantitative proteomics results should be controlled for their quality and comparability, before performing statistical tests and biological interpretation, to prevent drawing incorrect conclusions. Although some software platforms such as MaxQuant (3) incorporate normalisation as a step in data processing, one should always explore data distribution, consider variability of measurements and sample to sample correlation to control for biases or systematic errors. QC procedure implemented in ComplexBrowser consists of following parts:

- Box plot of log-transformed intensities
- Missing value bar plots
- Scatter plots with sample to sample correlation
- Histograms of CV
- Q-value charts
- Volcano plots
- PCA results

#### Data distribution – box plot and normalisation

Investigation of data distribution using box plots provides an easy way to spots differences in the signal intensities between two samples, that might be caused e. g. by injecting unequal amounts of peptide mixtures for the analysis. Note that box-plots use log-transformed values, hence if the whole range of values is shifted by only 1 unit on y axis (looking at median), it means that median value of in one sample is actually 2 fold greater than in the other. In such case normalisation is recommended.

ComplexBrowser implements four normalisation methods based on Normalyzer (4):

1. Mean normalisation – mean total intensity for each sample is calculated, then measurements within each sample are divided by their mean (equalizing mean of all samples to 1). Finally the intensities in each sample are calculated by multiplying each column by average total intensity of the original dataset.
2. Median – median total intensity for each sample is calculated, then measurements within each sample are divided by their median (equalizing median of all samples to 1). Finally the intensities in each sample are calculated by multiplying each column by median total intensity of the original dataset.
3. Total intensity – mean total intensity for each sample is calculated, then measurements within each sample are divided by their sum (expressing intensities as a fraction of total signal). Finally the intensities in each sample are calculated by multiplying each column by average total intensity of the original dataset.
4. Quantile normalisation aims to force sets to have perfectly aligned distribution. Since we consider this procedure may minimize experimental differences, ComplexBrowser applies quantile normalisation only within samples from the same conditions and later uses total intensity method.

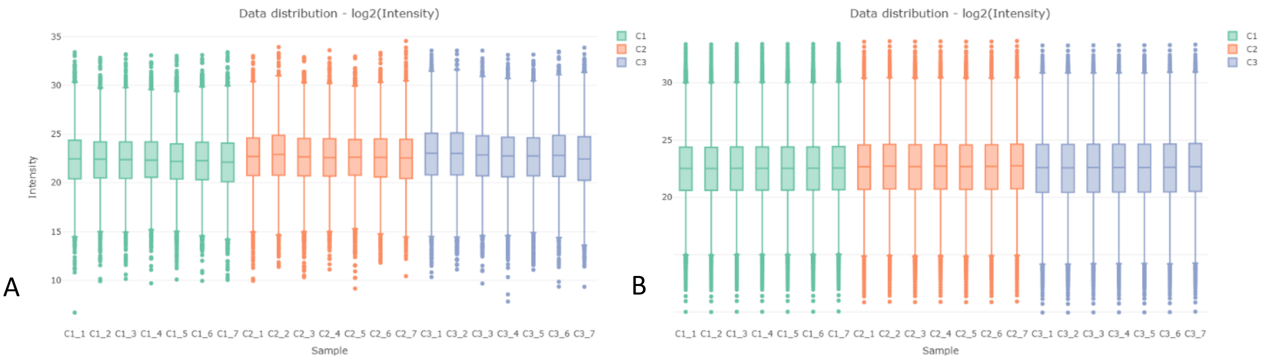


Figure 1. A - An example dataset before normalization, visible variation in medians. B - The same dataset after quantile normalisation with equalised medians and very similar, comparable distributions.

#### Missing values

Experiments with label-free quantitation deal with the problem of missing values, as due to large number of co-eluting peptides and stochastic selection of ions for fragmentation, inconsistencies in peptide identification and therefore quantification occur. This problem is partially solved by using spectra aligning algorithms that transfer peptide identification across samples (3). Nevertheless, data from analysis of complex samples rarely contains valid measurements for all proteins. Number of missing values in each injection can be an indicator of data quality and large variation in this parameter may point towards technical errors (e. g. with spray stability). ComplexBrowser delivers this information using a bar chart, where each bar corresponds to one sample.

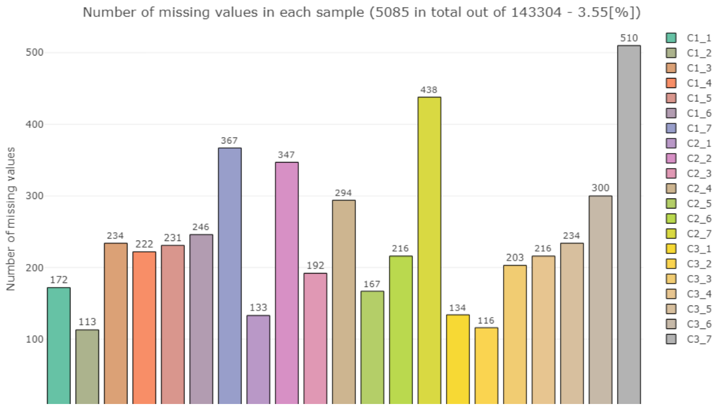


*Figure 2. Missing values in a sample dataset. Relatively large variation of MV number. Sample 3_7 with 510 measurements missing.*

#### Coefficient of variation distribution

Coefficient of variation (relative standard deviation) defined as: $\boldsymbol{c_v} = \frac{\sigma}{\mu}$ is calculated for absolute intensity measurements of one protein within a condition. Histogram of CV values for all proteins within given condition is plotted to estimate quantitation variability. Our observation is that a mean CV value <15% label-free, cell culture data indicates good quality. For TMT quantitation this value should be lower (5-10%).

#### Sample to sample correlation

Log-intensities of all proteins from one condition are plotted on the X axis and corresponding values from another condition on the Y axis. Pearson/Spearman/Kendal correlation is then calculated to assess run to run similarity. One should expect samples within the same biological condition to have a larger correlation between each other than to another condition.

#### Q-values

A simple line plot indicates the number of differentially expressed features based on a given false discovery rate threshold, helpful to estimate FDR cut-off for the further part of analysis.

##### Volcano plots

Well established plot type for visualising magnitude of effect and differentially expressed features. Values on X axis are log2(Ratios) for each protein, on Y axis – log10(FDR) values are plotted.

##### Principal component analysis

PCA is a dimensionality reduction procedure. If we consider samples as data points and quantitative measurements of all proteins as their coordinates, PCA aims to transform our dimensions to new find orthogonal coordinates that are able to highlight the differences between data points.

Samples from the same biological condition (replicates) should cluster together on a PCA plot, while different conditions should be separated.

#### Protein Complex Analysis

Additional parameters have to be set for analysis of protein complexes:

- q value threshold – **for visualization purposes only**
- Fold change threshold – for visualisation and summary purposes
- Noise threshold – for summary purposes
- Database (CORUM or EBI Complex Portal)
- Species the samples were generated from

Upon pressing "Run analysis" ComplexBrowser searches for relevant Uniprot accessions in user's data and displays all protein complexes with at least one subunit found in the input. For complexes with at least 3 quantified subunits complex fold change (CFC) is calculated. ComplexBrowser employs FARMS algorithm to determine complex fold changes.

FARMS (5) is based on Bayesian factor analysis with assumption of Gaussian measurement noise. It has previously been used in relation to protein inference and quantitation problem (6), and has proved to be useful in detecting outliers in peptide expression profiles, based on their covariance . In ComplexBrowser we employ it in an analogical approach. We use log-transformed subunit abundances across all samples to determine how coherently regulated they are. According to that, proteins are assigned a weight, by which their expression is scaled. Complex expression is then calculated in two steps.

First, scaled intensities of all subunits are summed within each replicate, subsequently the sums are averaged to give one complex expression value for each condition. Complex fold change is defined as relative change of expression between two conditions. ComplexBrowser also provides a noise level for each complex - a summary measure, describing variability in its expression profile, it takes a value between 0 (good) and 1 (bad). Recommended noise threshold is 0.5.

#### Complex table

Main table contains unique identifiers for each protein complex, its name, number of subunits, number of subunits found in the input, complex coverage, defined as the percentage of quantified subunits over all subunits forming the complex, subunits' Uniprot accessions, GO annotations, a PubMed reference, fold change values and the noise level.

Table 2: *Protein complexes found in the input dataset (using 1267 proteins)*

| No | ComplexID | Complex_Name | NUS | NQS | Coverage | Subunits | GO_terms | PubMedID | FC C2/C1 | FC C3/C1 | FC C4/C1 | FC C5/C1 | FC C6/C1 | Noise |
|---|---|---|---|---|---|---|---|---|---|---|---|---|---|---|
| | Al | All | . | . | . | A | All | All | A | A | A | A | A | A |
| 1 | 1 | BCL6-HDAC4 complex | 2 | 1 | 50 | P41182, P56524 | GO:0006265;GO:0... | 11929873 | | | | | | |
| 2 | 12 | BLOC-2 (biogenesis of lysosome... | 3 | 1 | 33.33 | Q86YV9, Q969F9,... | GO:0007032;GO:0... | 15030569 | | | | | | |
| 3 | 15 | NCOR complex | 6 | 1 | 16.67 | O15379, O60907,... | GO:0006265;GO:0... | 12628926 | | | | | | |
| 4 | 23 | BLOC-1 (biogenesis of lysosome... | 8 | 4 | 50 | O95295, P78537,... | None | 15102850 | -1.08 | -1.031 | 1.035 | 1.016 | 1.229 | 0.785 |
| 5 | 27 | Arp2/3 complex | 7 | 7 | 100 | O15143, O15144,... | None | 9359840 | -1.043 | -1.046 | -1.146 | -1.321 | -1.403 | 0.046 |
| 6 | 29 | PA28gamma complex | 1 | 1 | 100 | P61289 | GO:0043161;GO:0... | 9325261 | | | | | | |
| 7 | 30 | PA28 complex | 2 | 2 | 100 | Q06323, Q9UL46 | GO:0043161;GO:0... | 9325261 | | | | | | |
| 8 | 32 | PA700 complex | 20 | 20 | 100 | O00231, O00232,... | GO:0043161;GO:0... | 9148964 | -1.013 | -1.007 | 1.042 | 1.113 | 1.152 | 0.126 |
| 9 | 36 | AP1 adaptor complex | 8 | 4 | 50 | O43747, O75843,... | GO:0006886;GO:0... | 9733768 | 1.019 | -1.108 | -1.19 | -1.261 | -1.261 | 0.198 |
| 10 | 41 | Mi-2/NuRD-MTA2 complex | 5 | 1 | 20 | O94776, O95983,... | GO:0006265;GO:0... | 15454082 | | | | | | |

Showing 1 to 10 of 1,248 entries    Previous 1 2 3 4 5 … 125 Next

Figure 3. ComplexBrowser's table 2 with information about protein complexes found in the input dataset. Empty fold change cells due to less than 3 quantified subunits. NUS – Number of unique subunits; NQS – Number of quantified subunits; Coverage – percentage of quantified subunits in relation to all unique within one complex.

#### Complex star-graph

Protein complexes are visualised in form of a star graph. The central node represent the complex and the leaf nodes its subunits. The size and colour of the leaf nodes depends on their fold change. The size of a node increases with the absolute fold change, the colour indicates up (green) or down regulation (red), the nodes will only be coloured if the absolute fold change is larger than the set threshold. Nodes, which expression does not change are coloured in blue, whereas subunits that were not found in the input dataset are shown in grey. There are two types of edge thicknesses, thin nodes represent subunits that were not differentially expresses, thick edges indicate that the q value for a given condition was below the set cut-off.

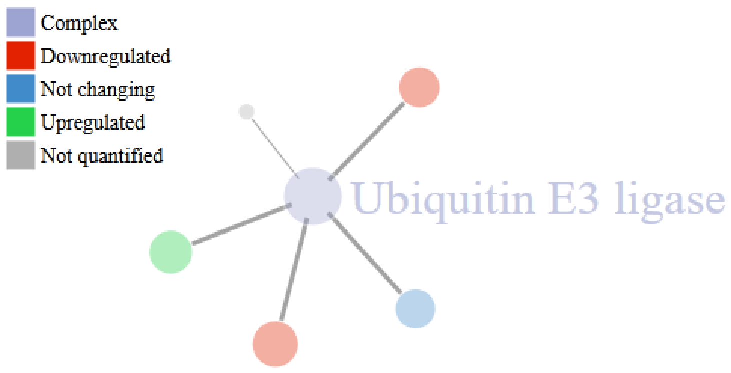

Figure 4. Complex star graph based on ubiquitin E3 ligase complex in condition 4 of the sample dataset. Thick edges represent proteins recognised as differentially expressed according to the set threshold.

#### Complex expression profile

To provide a more specific picture of various protein expression among complex's subunits a multiline plot of log intensities is drawn. This enables the user to quickly visually inspect trends in protein expression and the quality of correlation. By default plot is based on mean-normalised log-transformed protein abundances. Additionally, to account for measurement variability, the scale of the graph can be changed to Z-score based on log-values.

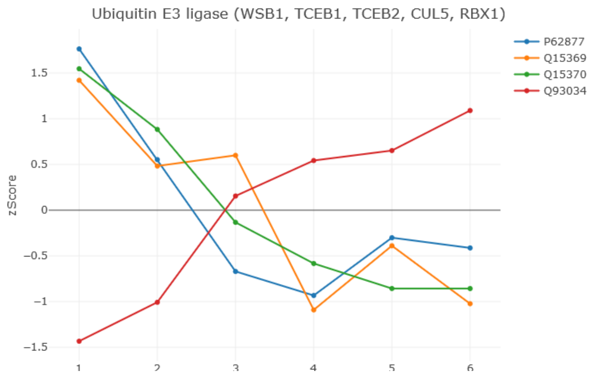

Figure 5. Ubiquitin E3 ligase expression profile. Protein Q93034 is a clear outlier, not following the expression trend spotted. (Seen in the previous section in star graph as green node)

#### Subunit expression plot

Bar plot of absolute intensities for a protein is drawn when a node is clicked on the star graph, allowing the user to investigate changes as well as standard deviations of protein abundance. This proves useful to to take a closer look at outliers spotted in the multiline plot.

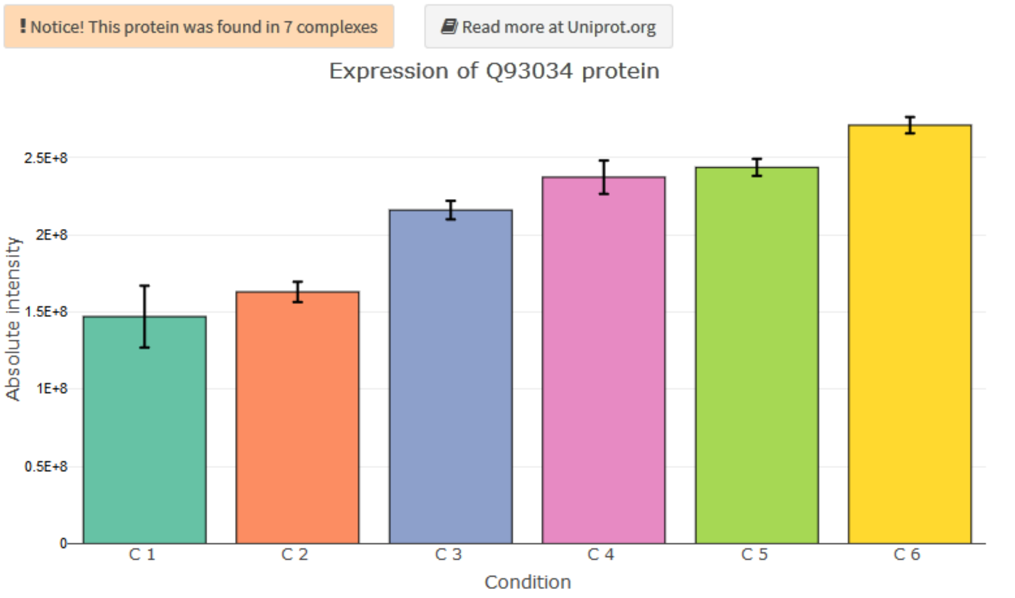


| | Fold change C2/C1 | Fold change C3/C1 | Fold change C4/C1 | Fold change C5/C1 | Fold change C6/C1 |
|---|---|---|---|---|---|
| Q93034 | 1.109 | 1.469 | 1.615 | 1.658 | 1.844 |

| | qValue C2 | qValue C3 | qValue C4 | qValue C5 | qValue C6 |
|---|---|---|---|---|---|
| Q93034 | 0.1688 | 0.0003 | 0 | 0 | 0 |

Figure 6. Expression of the spotted outlier. Q93034 is significantly changing in condition 4 (FC = 1.615), it also participates in 7 different complexes. Further investigation of its biological role necessary.

#### Complex expression correlation

Linearity of complex members co-expression correlation is assessed by plotting log2-transformed intensities of subunits on a scatter plot and calculating a correlation coefficient. Since both independent and dependent variables are affected by measurement technical variability we used orthogonal distance regression to calculate best model. A single $R^2$ value is returned as an indicator of co-expression linearity. As a rule of thumb, value > 0.95 can be considered good and >0.9 as acceptable.

#### Complex information

Additional complex information such as gene names for subunits, their full names, FunCat annotation, comment and associated disease are displayed for CORUM databse.

In case of EBI Complex Portal, ComplexBrowser shows details concerning complex's subunit stoichiometry is (if available), GO annotations, comment section and disease associations.

#### Expression heatmap

To display protein abundance changes and provide within complex clustering of proteins, a heatmap of log-transformed, mean normalized values is plotted. The hierarchical clustering included in the plot can be based on Manhattan, Euclidean, maximum or Minkowski distance with p = <3,20>. Clustering supports single, average, complete, mean, median and centroid linkage functions.

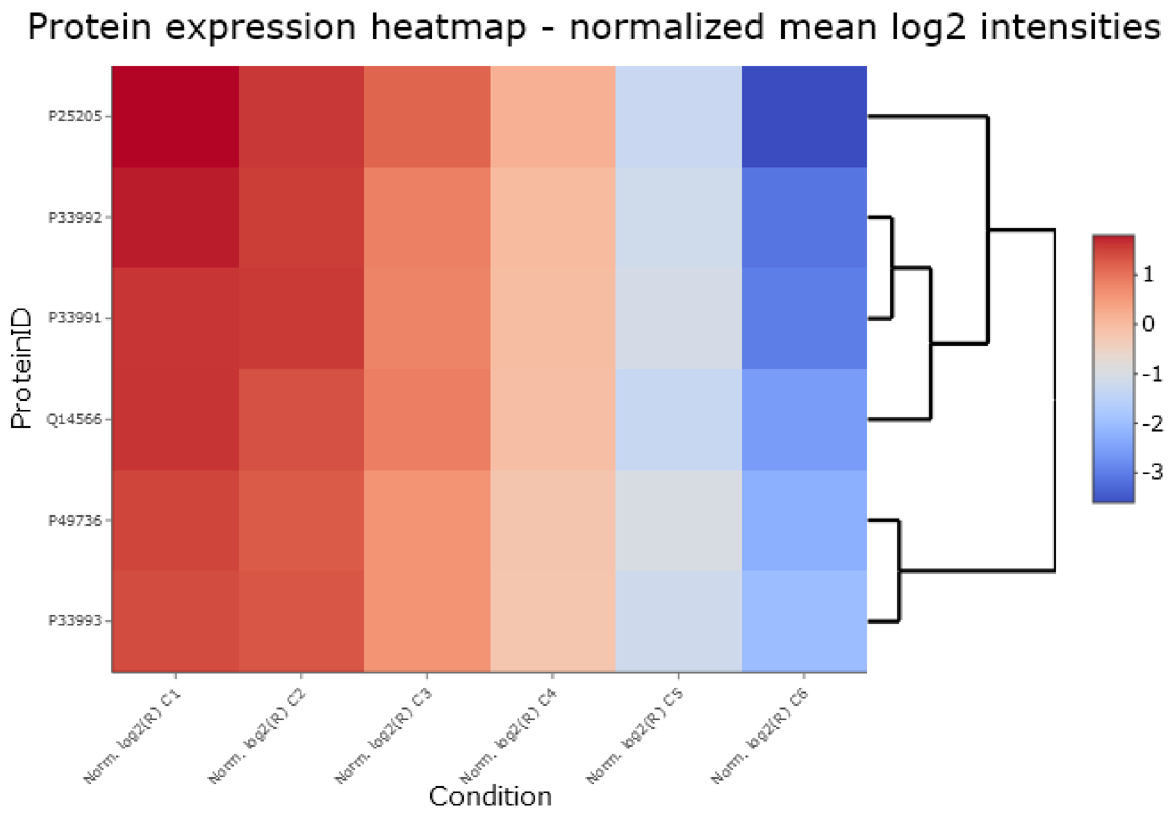


Figure 7. An example expression heatmap, based on well co-regulated MCM complex expression. Drawn using Euclidean distance and complete linkage function.

#### Expression correlation heatmap

Pairwise correlation (Pearson/Kendall/Spearman) of expression of complex subunits calculated based on intensities in all conditions is shown on the heatmap. This plot enables quick identification of subunits not following an expression trend, allowing the user to spot proteins that might take part in other quaternary structures. Dendrogram in correlation heatmap is based on the same distance measure and linkage function as selected for expression heatmap's hierarchical clustering.

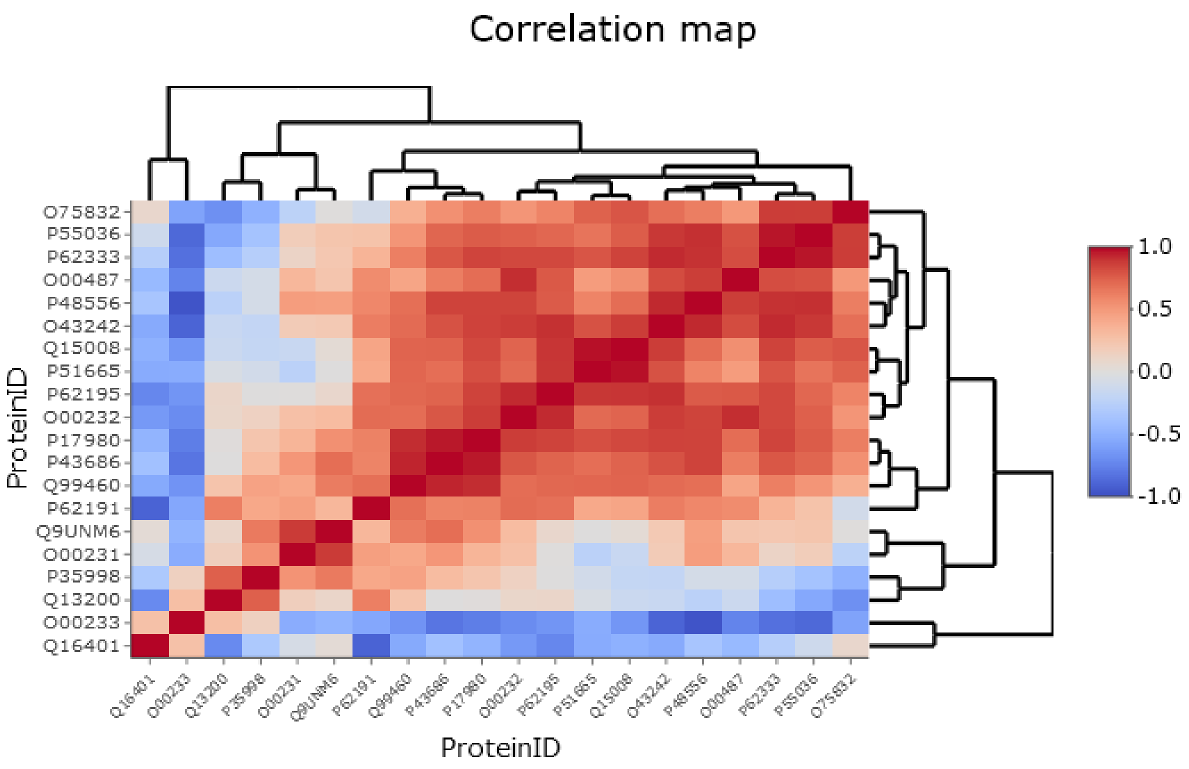


Figure 8. Based on co-expression of PA700 19S regulatory subunit of 26S proteasome across 6 conditions. Plot allows for identification of Q16401 and Q00233 proteins as outliers.

Both expression and correlation heatmaps are drawn for complexes with NQS higher or equal to 2. **Note** that looking at co-expression heatmap is informative only if comparing more than two conditions.

#### Summary

ComplexBrowser provides a summary bar plot, showing the number of regulated protein complexes in each condition. This plot takes into account threshold for noise and fold change set at the beginning of the analysis.

Additionally for each condition tables of 5 most upregulated and 5 most downregulated proteins are provided. Together this allows for quick assessment of global impact of the experimental condition on the molecular processes.

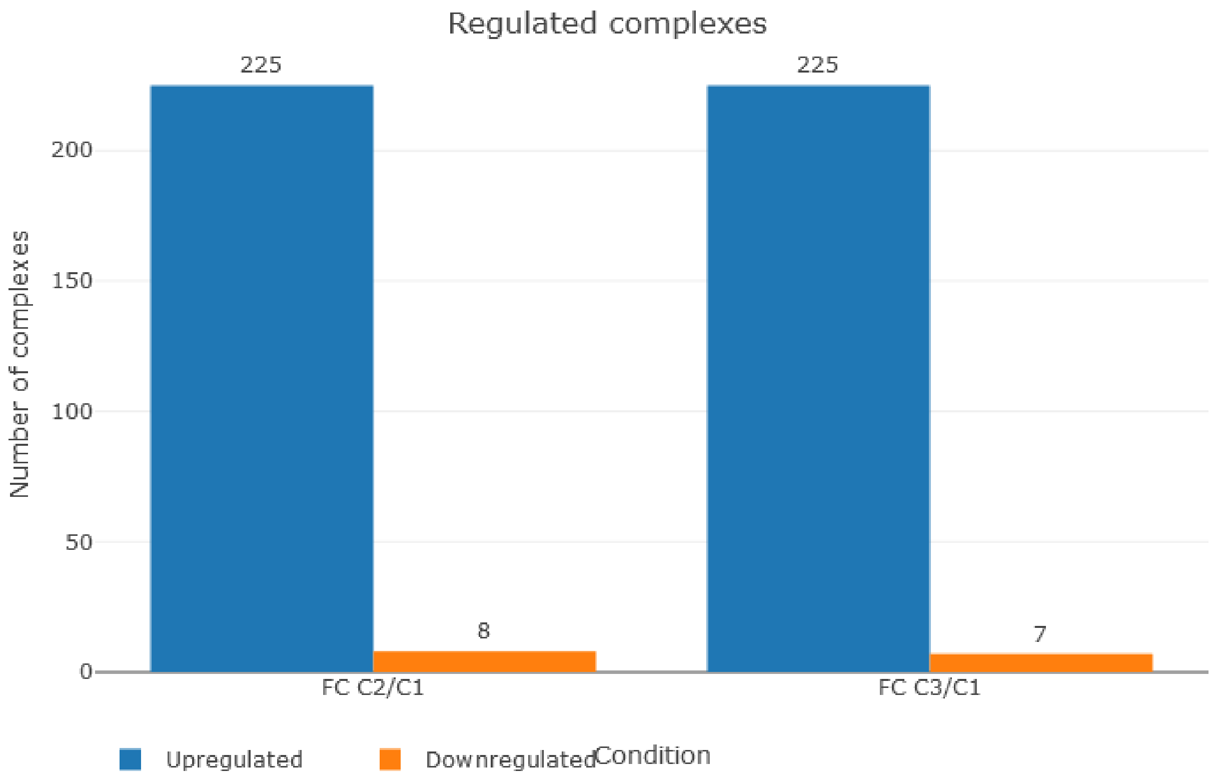


*Figure 9. Summary bar plot. Based on Adenocarcinoma data set from Wisniewski et al.* (7). *Thresholds set to 0.5 for noise and 1.5 for fold change.*
