## Supplementary material for "ComplexBrowser: a tool for identification and quantification of protein complexes in large scale proteomics datasets": File S2_QCreport_Adenocarcinoma

### ComplexBrowser - QC Report

This is a quality control generated on the 2019-02-28 by the ComplexBrowser. The input data had 3 conditions and 7 replicates with 6824 proteins quantified. We have found 5085 missing values in your data, that account to 3.548 [%] of all values.

#### Data characteristics

##### Distribution and correlation

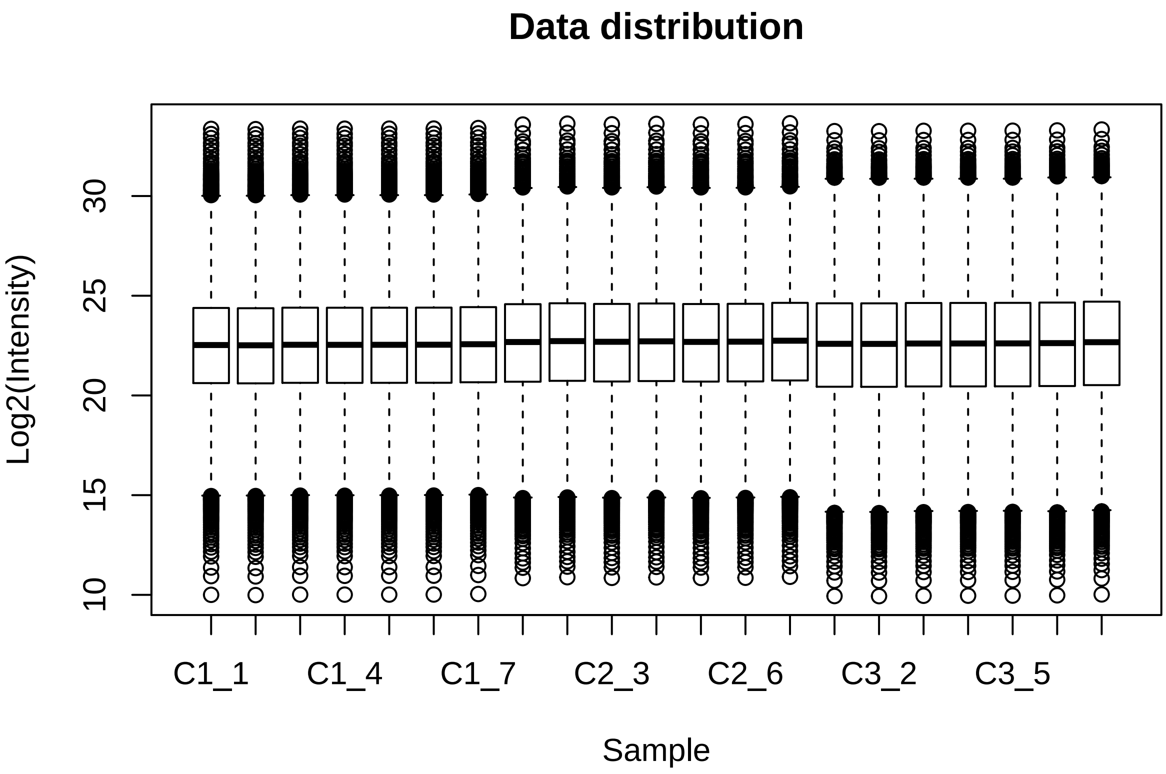


| | Cond. 1 | Cond. 2 | Cond. 3 |
|---|---|---|---|
| Min | 13.364 | 13.159 | 13.201 |
| Mean | 22.807 | 23.053 | 22.900 |
| Median | 22.627 | 22.889 | 22.764 |
| Max | 33.287 | 33.336 | 33.056 |

Table 1: Mean Log2(Intensity) values table

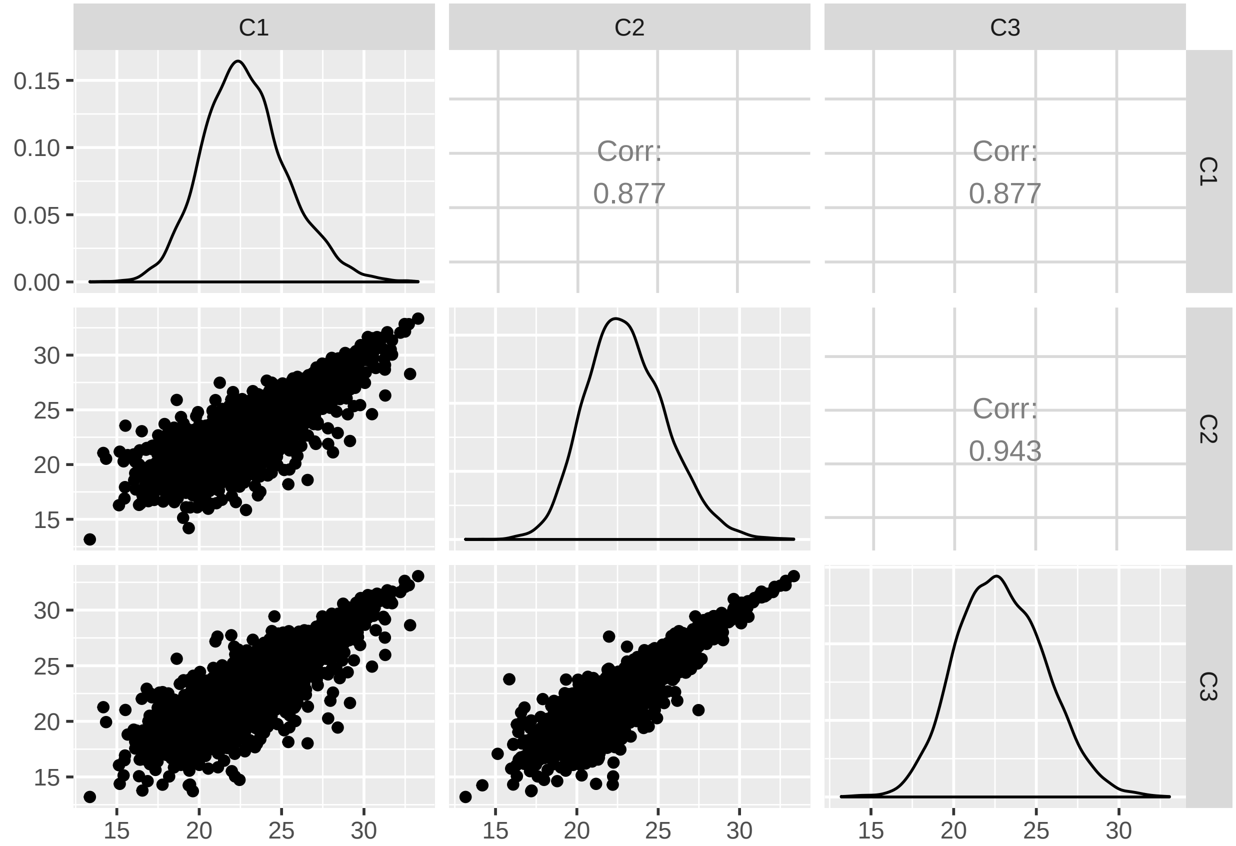
C1
C2
C3
0.15
0.10
0.05
0.00
Corr:
0.877
Corr:
0.877
C1
30
25
20
15
Corr:
0.943
C2
30
25
20
15
C3
15
20
25
30
15
20
25
30
15
20
25
30

**Missing values across samples**

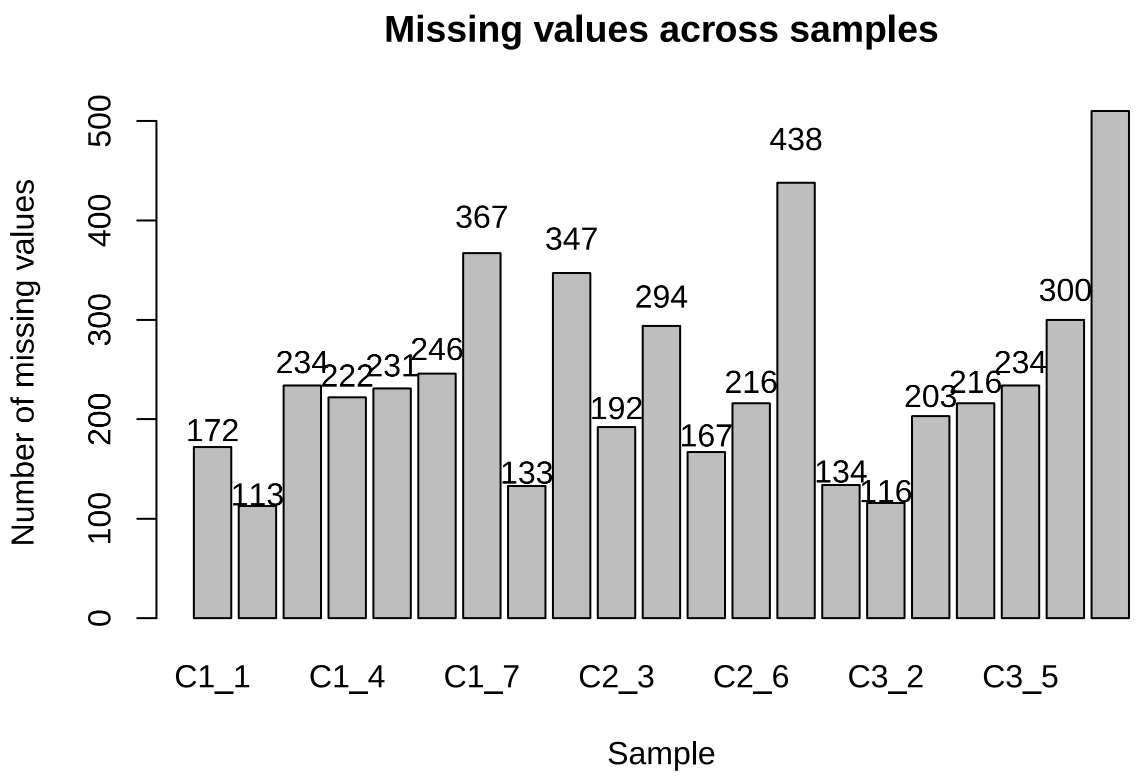

**Data quality - Coefficient of Variation**

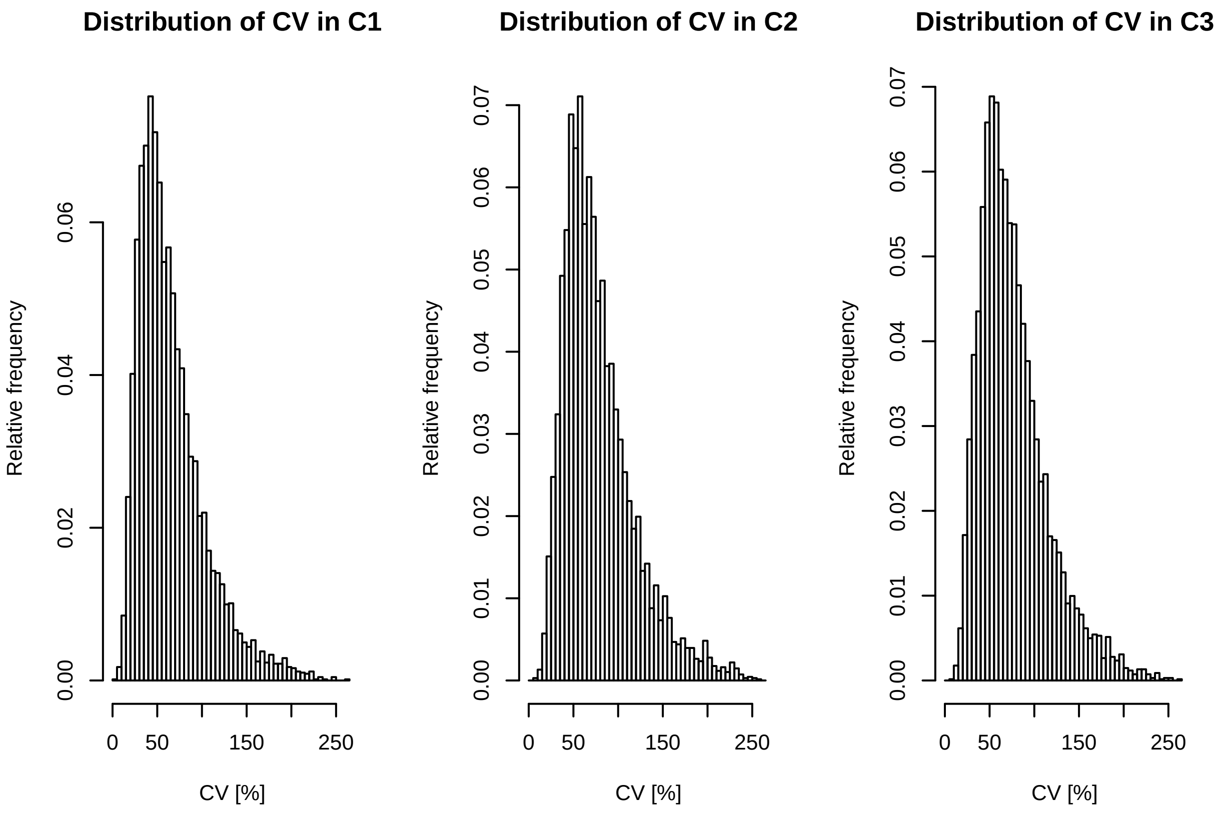


| | Cond. 1 | Cond. 2 | Cond. 3 |
|---|---|---|---|
| Min | 4.760 | 7.051 | 8.827 |
| Mean | 65.432 | 78.401 | 76.703 |
| Median | 56.415 | 69.635 | 69.015 |
| Max | 262.500 | 255.400 | 263.200 |

Table 2: Coefficient of Variation across conditions

In case of label free experiments median CV values should not exceed 15% for the data to be considered of good quality. For labeled experiments this value should be even lower.

**Data insights - Q Values and Fold Changes** Fold changes are calculated as:
$R = I_2/I_1$
$FC = R\ if\ R >= 1\ and\ FC = -1/R, for\ R < 1$

| | qValue C2/C1 | qValue C3/C1 |
|---|---|---|
| Min | 0.0000 | 0.0000 |
| Mean | 0.2921 | 0.2792 |
| Median | 0.2998 | 0.2764 |
| Max | 0.5940 | 0.5767 |

Table 3: Q Values across conditions

| | FC C2/C1 | FC C3/C1 |
|---|---|---|
| Min | -250.000 | -500.000 |
| Mean | 0.463 | -0.112 |
| Median | 1.236 | 1.136 |
| Max | 262.328 | 136.180 |

Table 4: Fold changes across conditions

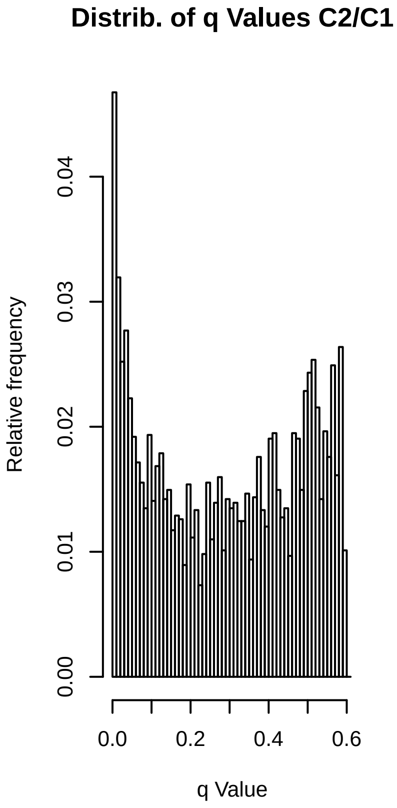
Distrib. of q Values C2/C1
Relative frequency
0.00
0.01
0.02
0.03
0.04
0.0
0.2
0.4
0.6
q Value

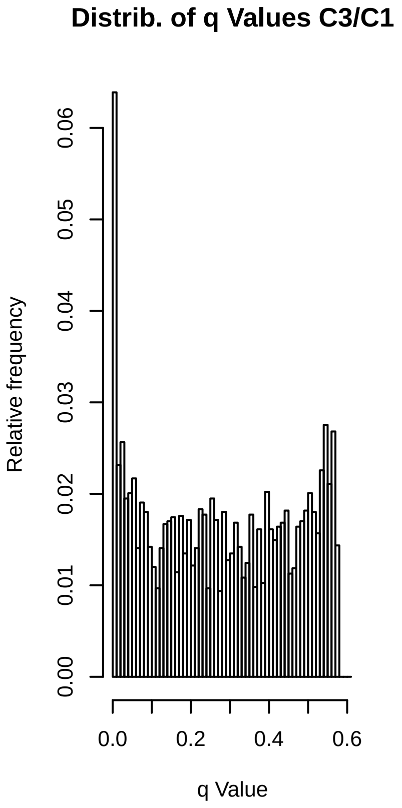
Distrib. of q Values C3/C1
Relative frequency
0.00
0.01
0.02
0.03
0.04
0.05
0.06
0.0
0.2
0.4
0.6
q Value

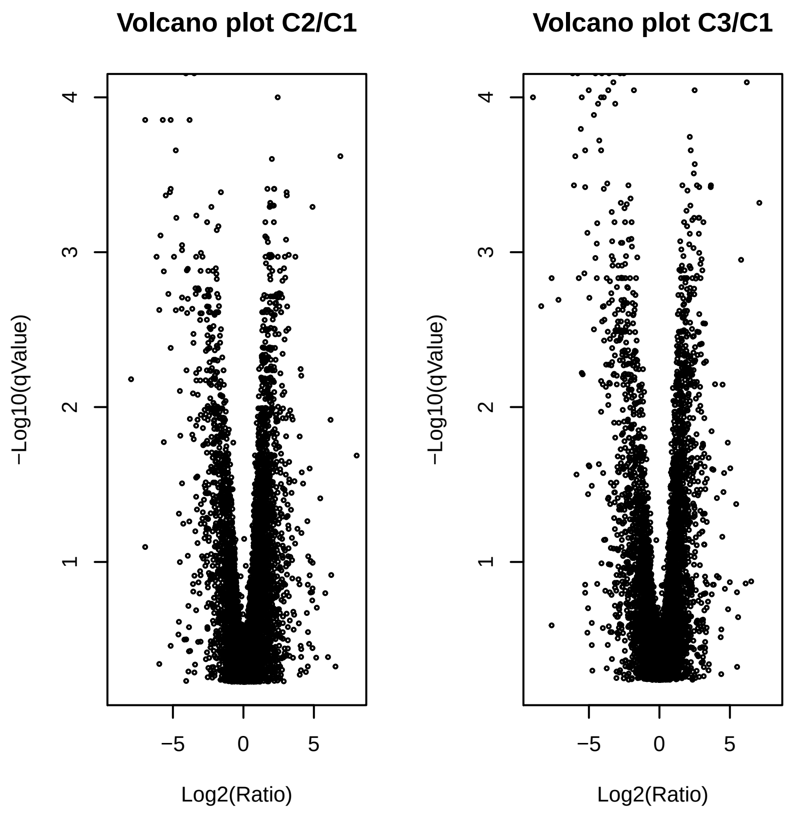
Volcano plot C2/C1
Volcano plot C3/C1
−Log10(qValue)
Log2(Ratio)
1
2
3
4
−5
0
5

**Data insigts - Principal Component Analysis**

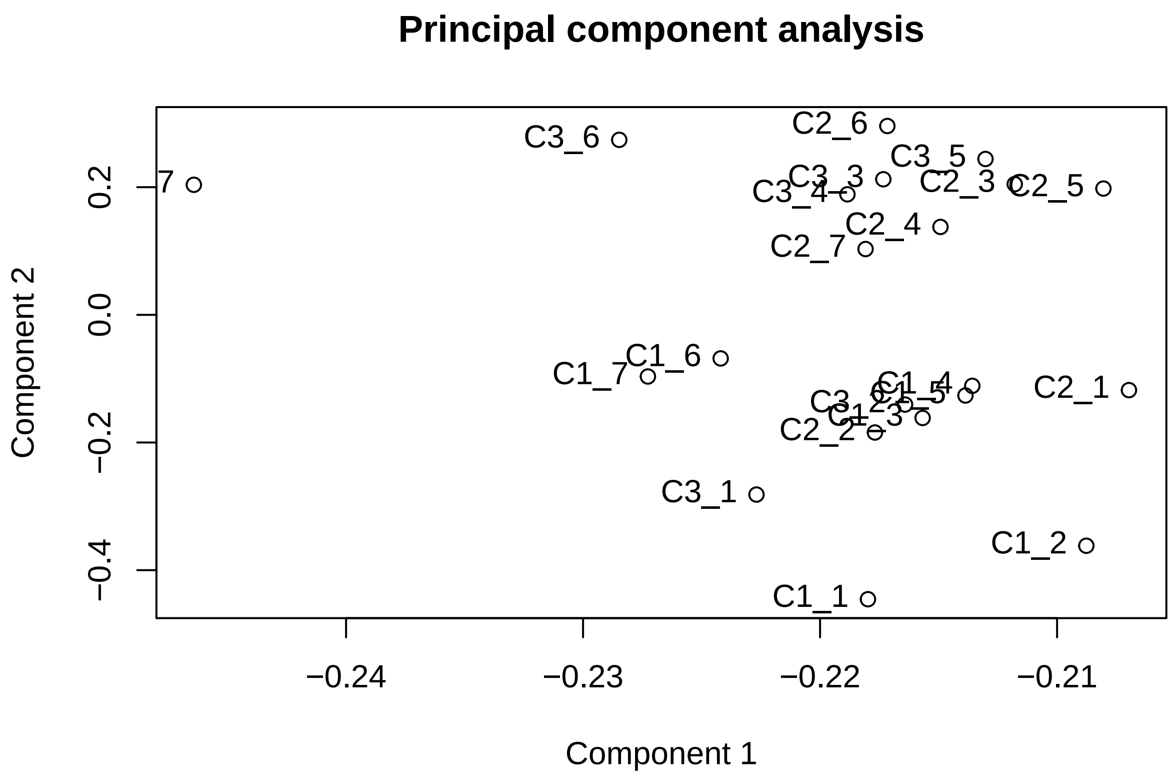
