## Supplementary material for "ComplexBrowser: a tool for identification and quantification of protein complexes in large scale proteomics datasets": File S3_QCreport_T-cell

### ComplexBrowser - QC Report

This is a quality control generated on the 2019-03-02 by the ComplexBrowser. The input data had 4 conditions and 2 replicates with 8431 proteins quantified. We have found 0 missing values in your data, that account to 0 [%] of all values.

#### Data characteristics

##### Distribution and correlation

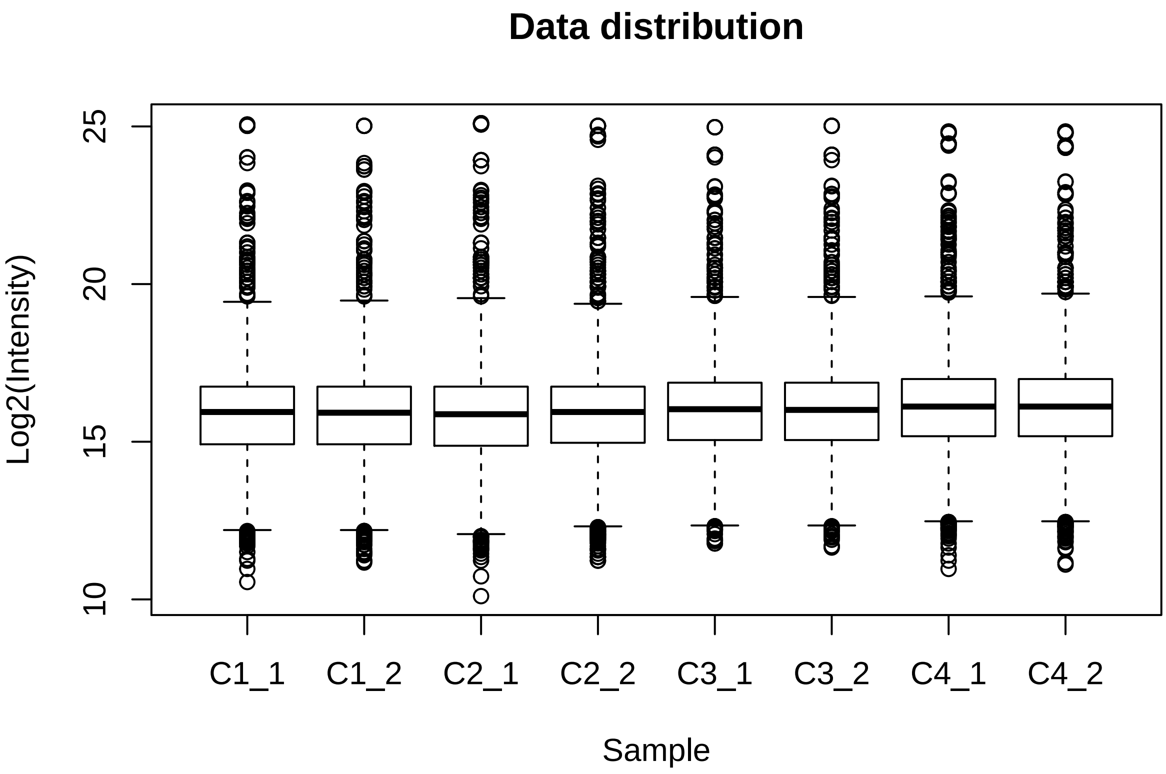


| | Cond. 1 | Cond. 2 | Cond. 3 | Cond. 4 |
|---|---|---|---|---|
| Min | 10.892 | 10.853 | 11.731 | 11.070 |
| Mean | 15.817 | 15.802 | 15.906 | 16.019 |
| Median | 15.932 | 15.920 | 16.021 | 16.116 |
| Max | 25.040 | 25.061 | 24.998 | 24.838 |

Table 1: Mean Log2(Intensity) values table

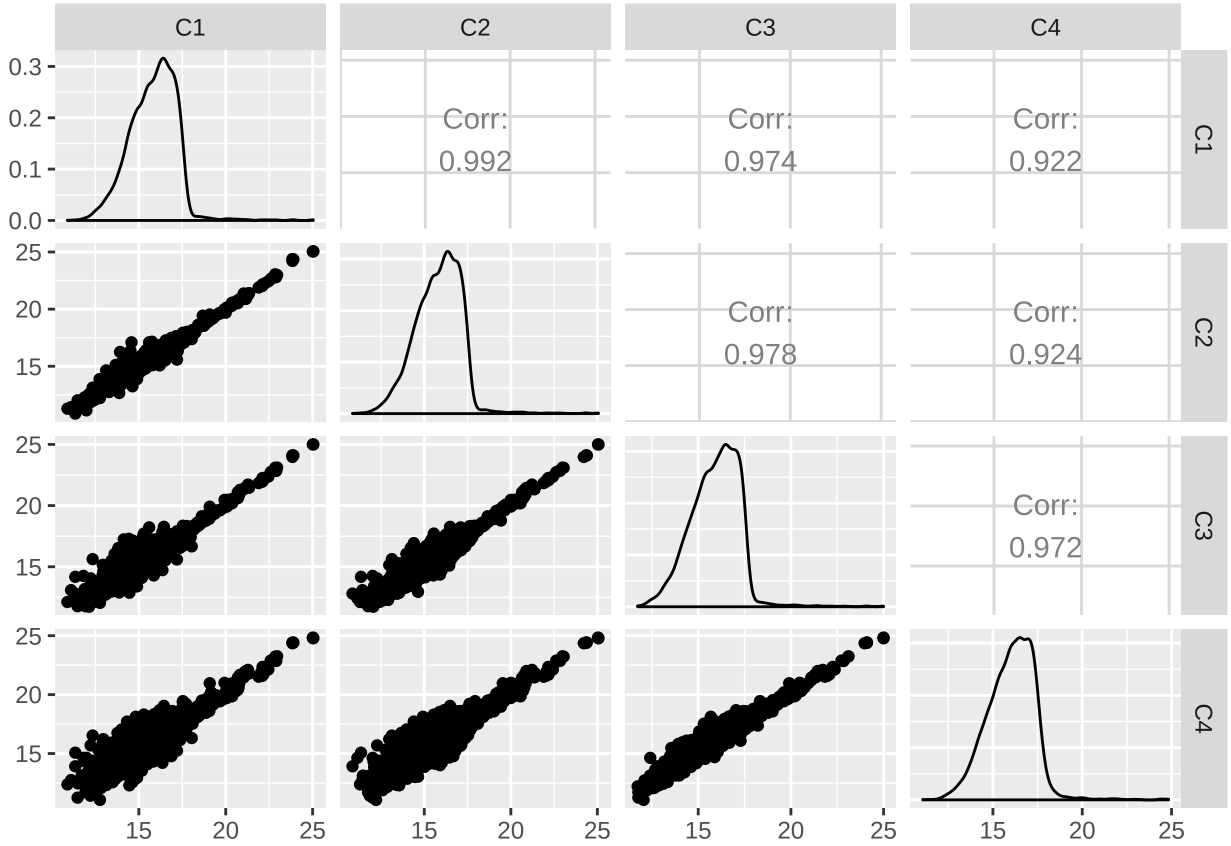
C1
C2
C3
C4
Corr:
0.992
Corr:
0.974
Corr:
0.922
Corr:
0.978
Corr:
0.924
Corr:
0.972
0.0
0.1
0.2
0.3
15
20
25

**Missing values across samples**

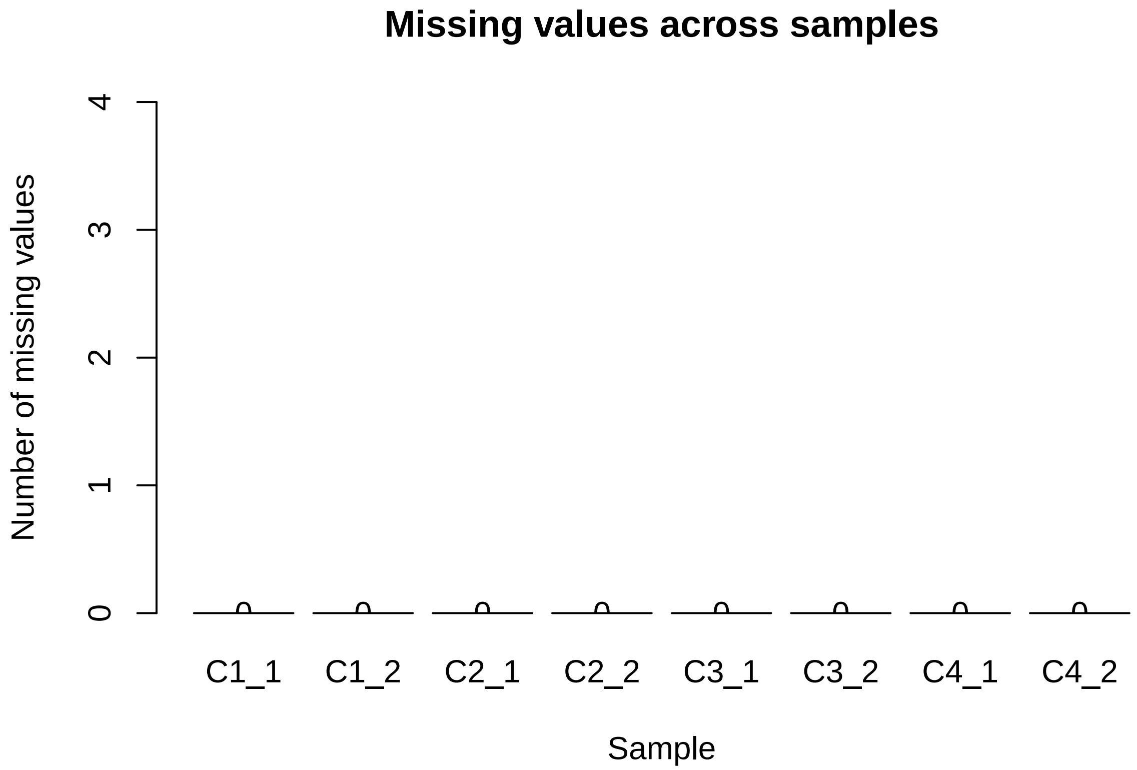

**Data quality - Coefficient of Variation**

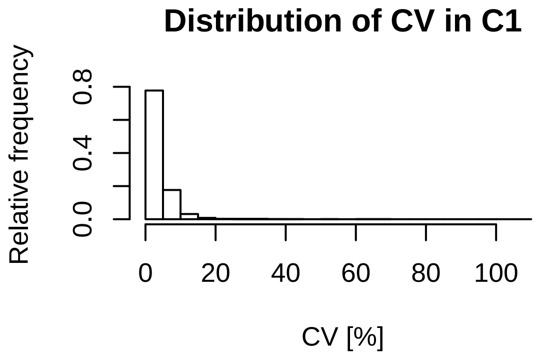


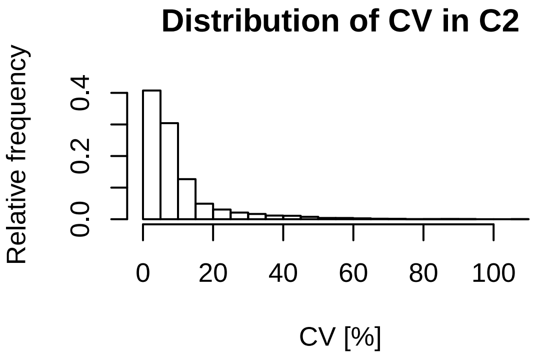


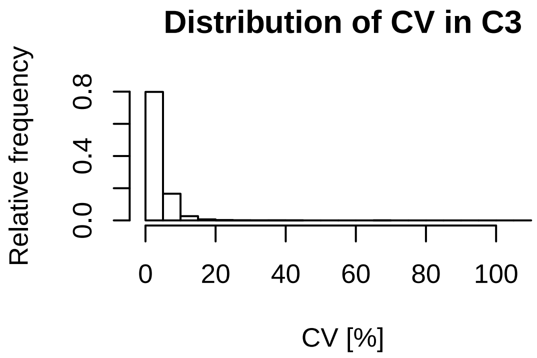


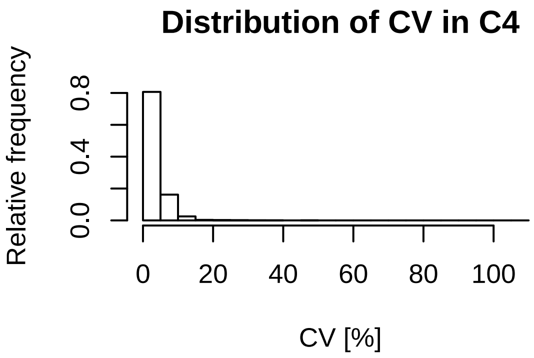


| | Cond. 1 | Cond. 2 | Cond. 3 | Cond. 4 |
|---|---|---|---|---|
| Min | 0.000 | 0.000 | 0.000 | 0.000 |
| Mean | 3.175 | 9.406 | 2.887 | 2.785 |
| Median | 2.245 | 6.149 | 2.111 | 2.050 |
| Max | 68.430 | 107.600 | 69.390 | 47.140 |

Table 2: Coefficient of Variation across conditions

In case of label free experiments median CV values should not exceed 15% for the data to be considered of good quality. For labeled experiments this value should be even lower.

**Data insights - Q Values and Fold Changes** Fold changes are calculated as:
$R = I_2/I_1$
$FC = R \; if \; R >= 1 \; and \; FC = -1/R, for \; R < 1$

| | qValue C2/C1 | qValue C3/C1 | qValue C4/C1 |
|---|---|---|---|
| Min | 0.0002 | 0.0004 | 0.0000 |
| Mean | 0.7200 | 0.3920 | 0.0815 |
| Median | 0.7048 | 0.3212 | 0.0093 |
| Max | 1.0000 | 1.0000 | 0.4632 |

Table 3: Q Values across conditions

| | FC C2/C1 | FC C3/C1 | FC C4/C1 |
|---|---|---|---|
| Min | -3.058 | -3.115 | -4.464 |
| Mean | 0.015 | 0.346 | 0.495 |
| Median | 1.000 | 1.027 | 1.079 |
| Max | 5.714 | 9.712 | 18.173 |

Table 4: Fold changes across conditions

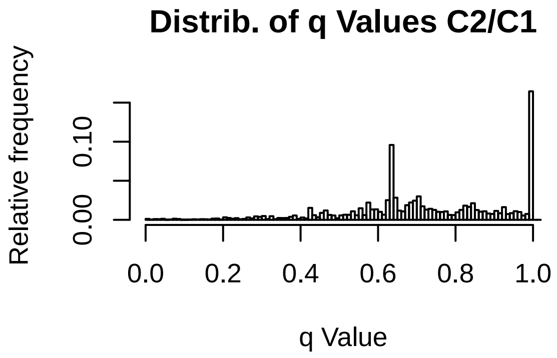
Distrib. of q Values C2/C1
Relative frequency
0.00
0.10
0.0
0.2
0.4
0.6
0.8
1.0
q Value

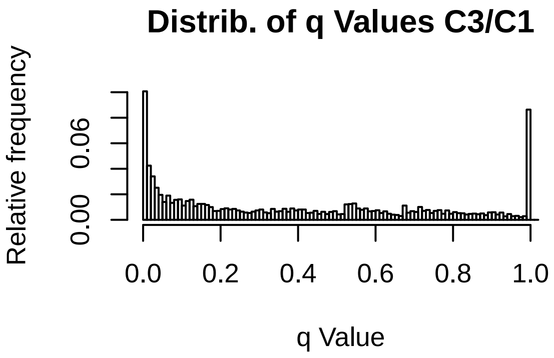
Distrib. of q Values C3/C1
Relative frequency
0.00
0.06
0.0
0.2
0.4
0.6
0.8
1.0
q Value

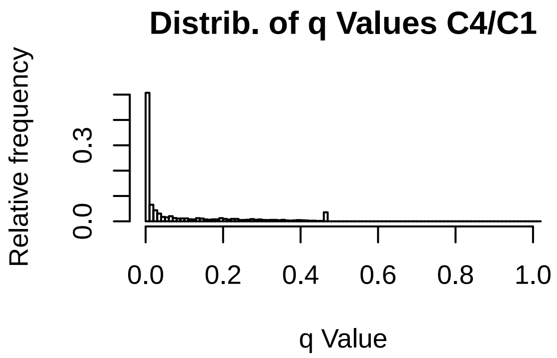
Distrib. of q Values C4/C1
Relative frequency
0.0
0.3
0.0
0.2
0.4
0.6
0.8
1.0
q Value

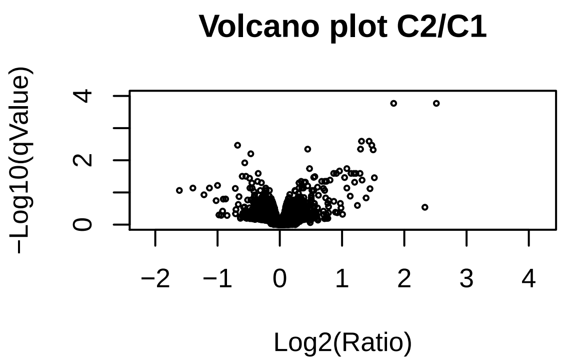
Volcano plot C2/C1
−Log10(qValue)
Log2(Ratio)
0
2
4
−2
−1
0
1
2
3
4

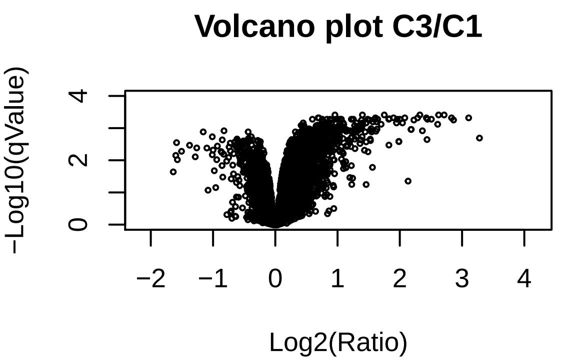
Volcano plot C3/C1
−Log10(qValue)
Log2(Ratio)
0
2
4
−2
−1
0
1
2
3
4

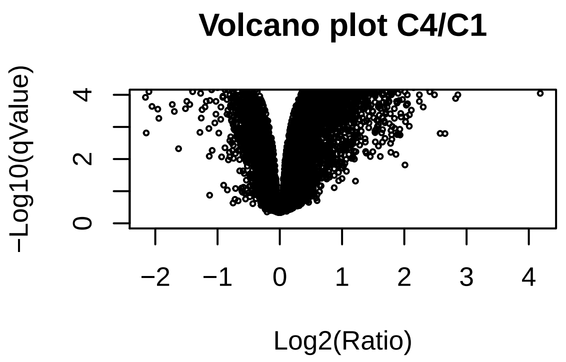
Volcano plot C4/C1
−Log10(qValue)
Log2(Ratio)
0
2
4
−2
−1
0
1
2
3
4

**Data insigts - Principal Component Analysis**

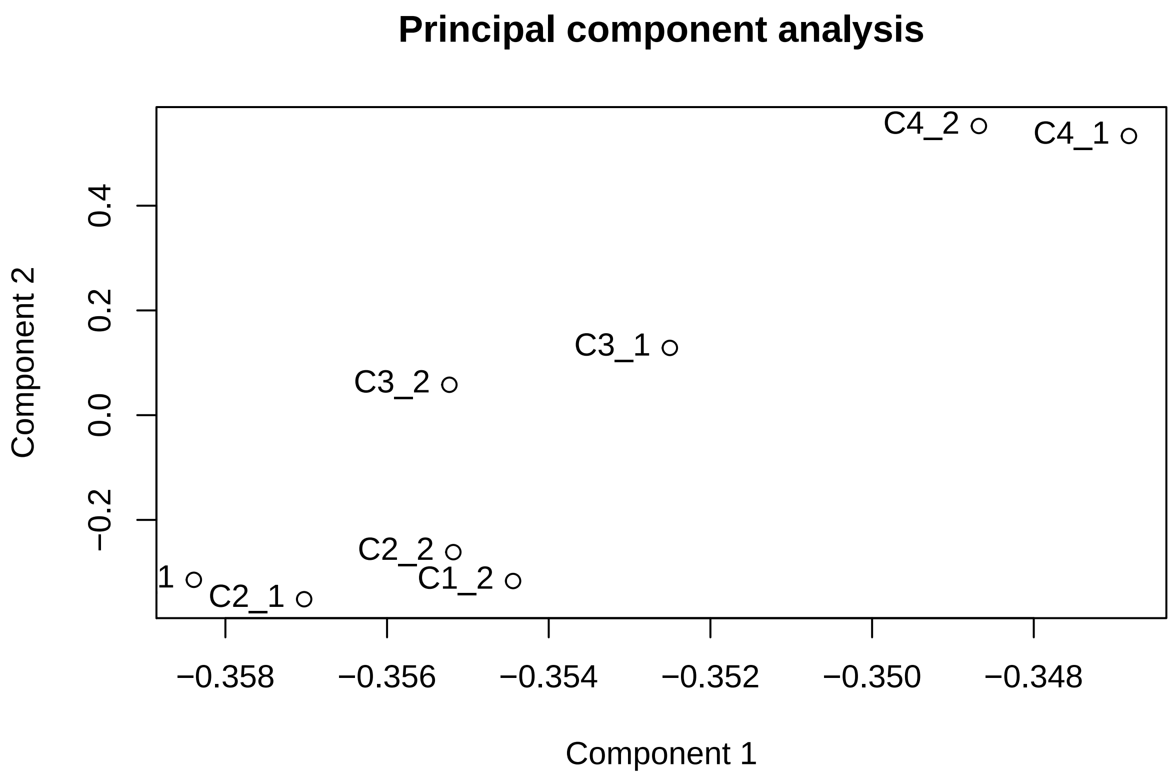
